# An amphipathic helix drives interaction of Fibrillins with plastoglobule lipid droplets

**DOI:** 10.1101/2023.09.28.559984

**Authors:** Kiran-Kumar Shivaiah, Duncan M. Boren, Andres Tequia-Herrera, Josh Vermaas, Peter Knut Lundquist

**Affiliations:** Department of Biochemistry and Molecular Biology, Michigan State University, East Lansing, MI 48824; Plant Resilience Institute, Michigan State University, East Lansing, MI 48824

**Author notes:** Department of Biological Chemistry, University of Michigan, Ann Arbor, MI 48109. Corresponding author: Peter Knut Lundquist.

**Keywords:** Plastoglobule, lipid droplet, Fibrillin, Plastoglobulin, amphipathic helix, lipids, lipocalin

## Abstract

Plastoglobule lipid droplets of chloroplasts serve complex roles affecting plant development, stress tolerance and photosynthesis. They harbor a set of approximately 42 proteins that collectively dictate plastoglobule functions. Due to the monolayer structure of plastoglobules which encompass a neutral lipid core, these proteins must associate monotopically on the plastoglobule surface. However, targeting determinants have not been identified for plastoglobule proteins, and the protein-membrane interaction mechanisms that establish the plastoglobule proteome remain unclear. Here, we demonstrate that plastoglobule-localized Fibrillins harbor an amphipathic helix at the lip of their β-barrel that is necessary for proper plastoglobule association. Molecular dynamics simulations support the specific interaction of the amphipathic helix of AtFBN1a with membranes rich in lipid packing defects which are expected to be especially prevalent on the tightly curved surface of plastoglobules. Introduction of one of the amphipathic helices into stromal-or thylakoid-localized FBNs was ineffective at redirecting the proteins to plastoglobules, likely due to endogenous protein-protein interactions that override the influence of the amphipathic helix. Proteomic analyses indicate AtFBN1a influences the plastoglobule proteome through outcompeting and recruiting specific proteins. We also demonstrate that the plastoglobule-localized FBNs, AtFBN1a and AtFBN7a, bind unsaturated fatty acids, particularly C18:1, and that elimination of the amphipathic helix suppresses fatty acid binding in AtFBN1a, but promotes fatty acid binding in AtFBN7a. Predicted amphipathic helices can be identified on two-thirds of plastoglobule proteins, indicating the use of amphipathic helices may be a general mechanism by which proteins selectively associate with plastoglobules.

## INTRODUCTION

Plastoglobules are lipid droplets (LDs) of plant chloroplasts, present in all photosynthetic organisms (1). As LDs, they are characterized by a monolayer polar lipid surface enclosing a neutral lipid core (2). Plastoglobules serve as a crossroads of metabolic activity within the chloroplast and appear to be essential for plant survival and stress resilience (1, 3–14). Based on mutant phenotypes and gene co-expression patterns, plastoglobules have been suggested to serve as regulatory and enzymatic hubs of chloroplast membrane remodeling and adaptation, crucial for plants to thrive within constantly fluctuating environments (1, 15).

Through the use of mass spectrometry-based proteomics, immunoblotting and confocal microscopy, a set of 42 proteins have been identified as *bona fide* plastoglobule-localized proteins in the model plant, *Arabidopsis thaliana* (7, 15–18). Consistent with the lipid-rich nature of plastoglobules, many of these proteins hold characterized or putative functions in lipid metabolism, including Tocopherol Cyclase, Carotenoid Cleavage Dioxygenase 4, and Allene Oxide Synthase. The most abundant proteins of the plastoglobule are seven members of the Fibrillin family, comprising approximately 50% of the *A. thaliana* plastoglobule protein mass (7, 15). FBNs share a lipocalin domain characterized by a β-barrel that forms a hydrophobic cavity capable of binding small hydrophobic ligands (19–21), and are believed to provide a structural role for the plastoglobule. Investigation of mutant lines in plastoglobule-targeted FBNs implicate them in stress tolerance (3, 4, 14, 22–24), as well as recruitment of proteins (such as the jasmonate biosynthetic pathway or carotenoid metabolism) to plastoglobules (3, 4, 13, 14, 25–28).

Despite the apparently essential nature of plastoglobules in photosynthetic organisms, the molecular mechanism(s) of protein targeting to plastoglobules remains unclear. Nearly all of the ca. 3000 chloroplast-localized proteins -including all 42 plastoglobule proteins - are nuclear encoded. N-terminally encoded chloroplast transit peptides (cTPs) direct these proteins to the Translocon at the Outer Chloroplast Envelope Membrane and Translocon at the Inner Chloroplast Envelope Membrane (TOC/TIC) protein import machinery which translocates the proteins across the chloroplast double envelope membrane. But once a plastoglobule protein has been imported, and cleaved of its cTP, how can it distinguish between the various membrane sub-compartments of the chloroplast?

Attempts to identify conserved sequence features among plastoglobule-localized proteins, through sequence alignments or hidden Markov Models, have failed to identify common sequences. Confocal microscopy analysis of serially truncated AtFBN7a indicated nearly the entire polypeptide may be necessary for proper plastoglobule targeting (29, 30). In contrast, closely related isoforms of *Zea mays* Phytoene Synthase separately localize to either the plastoglobule or a dual localization on bilayer membranes and aqueous stroma, indicating subtle sequence differences may underlie plastoglobule association (31). Consistent, with the latter observation, a 19-residue hydrophobic region of rice Phytoene Synthase, OsPSY2, could direct cargo protein to plastoglobules when localized at the C-terminus (32).

Protein targeting to cytosolic LDs is better understood than that of plastoglobules and occurs through two broad targeting pathways: the first, known as the Class I (or ERTOLD) pathway, targets proteins to the endoplasmic reticulum membrane through a hydrophobic hairpin loop which then migrate laterally to physically connected LDs; in the second pathway, known as the Class II (or CYTOLD) pathway, proteins are directly localized to LDs from the cytosol, often through an amphipathic helix (AH) (33–35). Because plastoglobules are physically connected to the thylakoid membrane, the Class I targeting pathway, in which proteins would be targeted first to the thylakoid, before lateral migration to the plastoglobule, could be employed. Alternatively, proteins could employ a Class II targeting pathway in which AHs or lipid modifications (for example) drive proteins directly to associate with the plastoglobule.

The use of AHs in LD association is logical. The monolayer structure of LDs is expected to foster the formation of lipid packing defects, defined as gaps in the polar headgroup surface through which underlying neutral lipid becomes exposed to the aqueous solvent (36–38). The lipid packing defects favor interaction of AHs through hydrophobic interactions between the exposed neutral lipid and the hydrophobic face of the AH, as well as polar or charged interactions between the lipid headgroups or aqueous solvent and the hydrophilic face of the AH. In the context of cytosolic LDs, AHs typically harbor bulky hydrophobic side chains in their hydrophobic face that are critical for effective LD interaction, seemingly by promoting initial anchoring to the LD surface (34, 38–41). In addition, the positioning of uncharged polar residues near the interface between hydrophobic and hydrophilic faces is important for membrane curvature sensing in many AHs (36, 42). However, unlike LDs of the cytosol, which are composed of phospholipids, the monolayer of plastoglobules is comprised of galactolipids. Moreover, the lipid core of plastoglobules is rich in prenyl-lipids and their derivatives, such as plastoquinone and xanthophyll esters (7, 13, 43), in contrast to the lipid core of cytosolic LDs which is characterized by triacylglycerol and sterol esters (1, 44, 45). Plastoglobules are also substantially smaller in size than cytosolic LDs, often as small as ca. 70 nm in diameter (10). Thus, it is not clear to what extent targeting principles that apply to cytosolic LDs can be translated to plastoglobules.

Here, we report that FBNs employ a unique AH insert within their lipocalin domain that drives effective plastoglobule targeting. Circular dichroism spectroscopy indicates a stable helical structure in the AH while molecular dynamics (MD) simulations support interaction of the AH sequence specifically with membrane surfaces enriched in lipid packing defects. We also demonstrate that FBNs bind fatty acids and that the efficacy and ligand preference is affected by the presence/absence of the AH insert. Predicted AHs are enriched in plastoglobule-localized proteins – found on well over half of all plastoglobule proteins – indicating the use of AHs may represent a general mechanism of plastoglobule targeting.

## RESULTS

### Predicted AHs are a common feature of plastoglobule proteins

To test whether AHs may be involved in association of proteins to plastoglobules we used the HeliQuest web-based software tool (46) to screen for predicted AHs amongst *A. thaliana* proteins that have been demonstrated to localize to plastoglobules in various proteomic experiments (7, 15, 16, 47). A total of forty-two plastoglobule proteins were screened for AHs of alpha-helical or 3-11 helical structure. HeliQuest allows customization of multiple search parameters, including the net charge state, the hydrophobic moment, and the maximal number of charged residues, among others. To first optimize the parameter settings, we ran the screen with six proteins known to contain AHs (*Homo sapiens* ADP-ribosylation Factor GTPase-activating enzyme [HsArfGAP] (48); *H. sapiens* Golgi Microtubule-associated Protein of 210 kDa [HsGMAP-210] (49); *H. sapiens* Perilipin 1, 3, and 4 [HsPLN1,3,4] (50–52), and *Penicillium chrysogenum* Peroxin-11 [PcPEX11] (53)), as well as two soluble stromal chloroplast proteins which are not expected to hold AHs (*A. thaliana* Caseinolytic Protease R1 [AtClpR1] (54), and *Chlamydomonas reinhardtii* Thioredoxin H [CrTrxH] (55)), serving as positive and negative controls, respectively. While multiple AHs were predicted within each of the positive controls, no AHs were predicted from CrTrxH and only a single abnormally short (11 residue) predicted AH was found in AtClpR1 (**Table S1**). This suggests that with the selected search parameters we were able to effectively identify AHs with a low false positive identification rate.

The HeliQuest search predicted at least one AH in 31 of the 42 plastoglobule proteins, including all six of the ABC1 kinases and all seven of the FBNs (**Table 1**), which represent the most abundant components of the plastoglobule proteome (7, 15). Indeed, predicted AHs were especially extensive among these two protein families, though multiple AHs were predicted in over half of all plastoglobule proteins (**Supp Data File**). Using the same HeliQuest parameters, we found that predicted AHs were especially prevalent among plastoglobule-localized proteins compared to those localized to the other chloroplast membrane compartments, the thylakoid and envelope (**Fig. S1**). Plastoglobule proteins lacking predicted AHs tended to be lower abundant members of the plastoglobule proteome (*e.g.* the UbiE-like Methyl Transferases and Flavin Reductase-like proteins), or those recruited to the plastoglobule under unique conditions (*e.g.* the Allene Oxide Cyclase isomers).

**Table 1.**
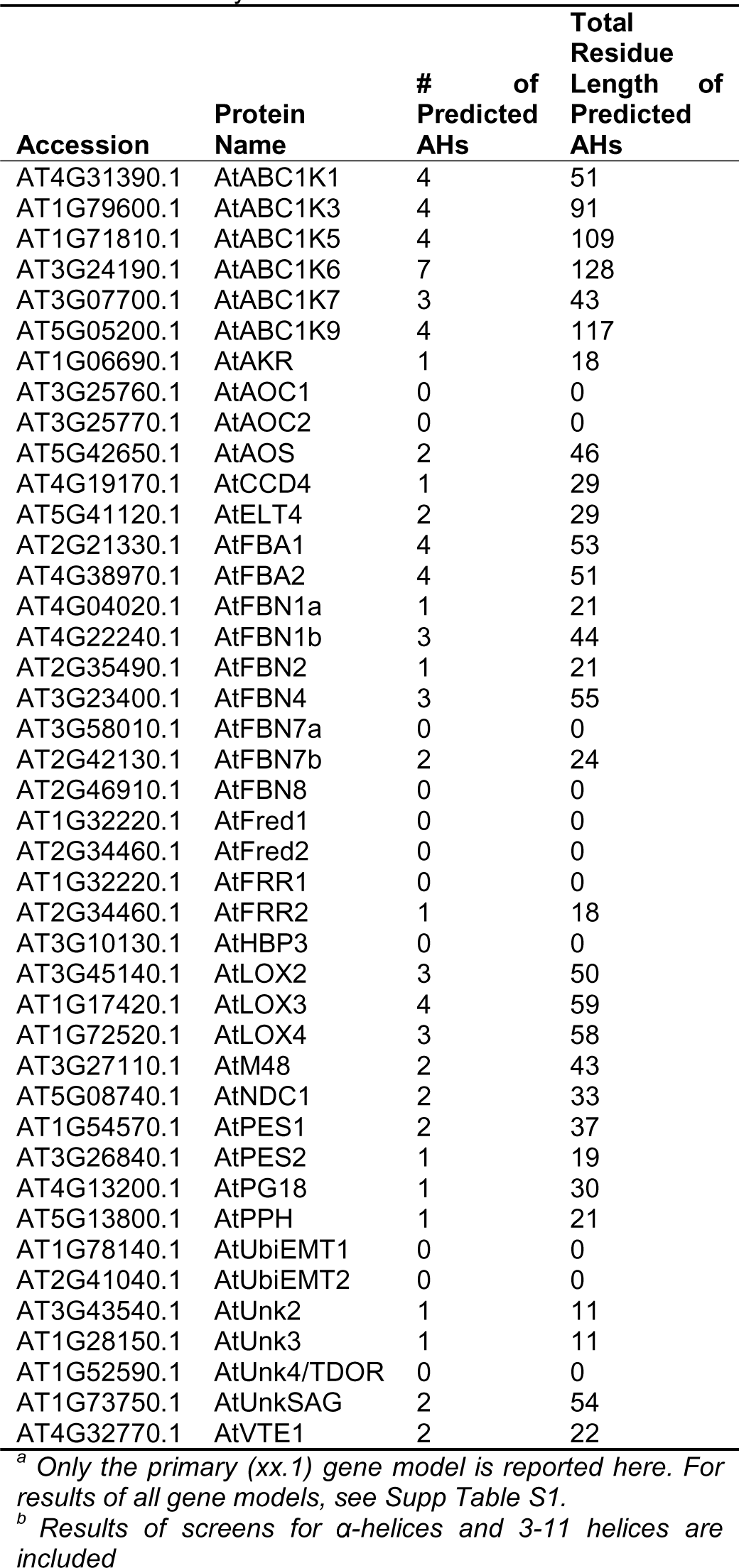
Predicted AHs Among Plastoglobule Proteins from a HeliQuest Survey^a,b^.

**Table 2.**
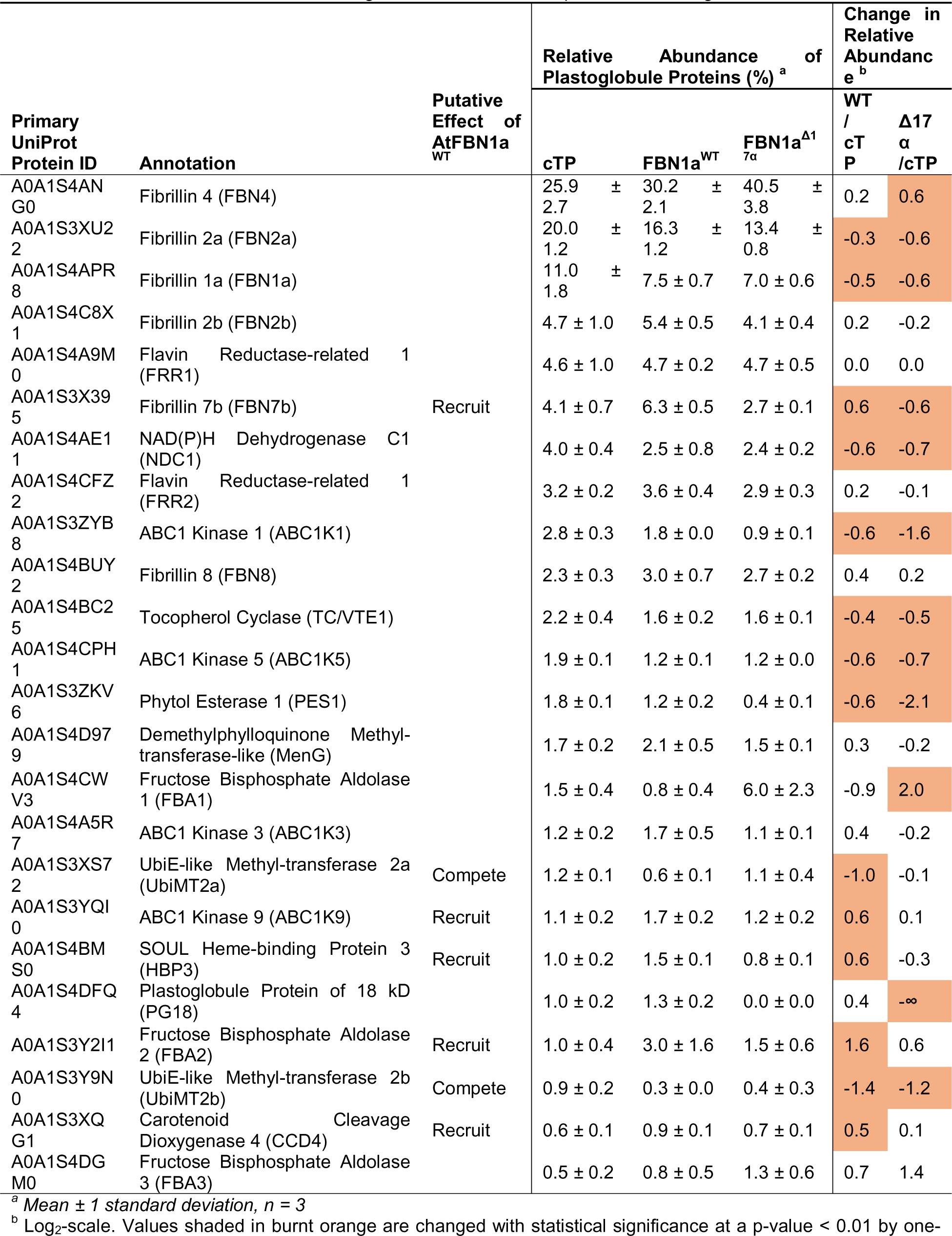
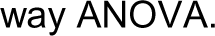
The *Nicotiana benthamiana* Plastogobule Proteome and Impacts of Heterologous Protein Accumulation.

### FBN Proteins of the Plastoglobule Harbor a Unique Helical Insert Sequence

To better understand their structural context, we highlighted the predicted AHs of each of the thirteen FBNs encoded in the *A. thaliana* genome within the context of structural models generated using AlphaFold (56, 57) and RoseTTAFold (58) (**Fig. S2**). Models from the two programs were highly similar and thus we relied on the AlphaFold models for our analyses. All FBN proteins were comprised of an eight-stranded β-barrel structure with a conserved GxW motif positioned near the bottom of the β-barrel, the Trp side-chain facing inside the barrel. This modeled structure is consistent with crystal structures of other lipocalin domains such as *Escherichia coli* Blc (59, 60). Upstream of the β-barrel (*i.e.* near the N-terminus) was found a crisscrossing α-helical pair of varying length. While the helical pair and β-barrel formed the shared core features of all thirteen *A. thaliana* FBNs, a unique helical insert was additionally found within all but one of the plastoglobule-localized FBNs, and was absent from all but one of the thylakoid/stromal-localized FBNs (**Fig. 1A**). This helical insert occurred between the fifth and sixth strands of the β-barrel (strands E and F according to the lipocalin domain nomenclature), spanned 16 to 27 residues in length, and was usually flanked by short β-strand inserts not present in the core FBN structure (**Fig. 1C, D**). The helical insert was also sometimes found in a kinked conformation, with an interruption to the helical structure of two or more residues (**Fig 1E**). Notably, temperature-induced lipocalin proteins (TILs) use a hydrophobic, proline-rich insert sequence in this same location, between β-strands E and F, for peripheral membrane association at the plasma membrane (61). Furthermore, the placement of the FBN helical insert along the side of the β-barrel opening was reminiscent of the cryo-EM structures of the human and fly Seipin lumenal domains, in which a hydrophobic helix (*i.e.* the ‘central helix’ or ‘membrane-anchored helix’) lies near the opening of an eight-stranded β-sandwich (**Fig. S3**). The hydrophobic helix of Seipin embeds within the endoplasmic reticulum membrane and is important for its oligomerization, function, and localization (62–64).

**Figure 1.**
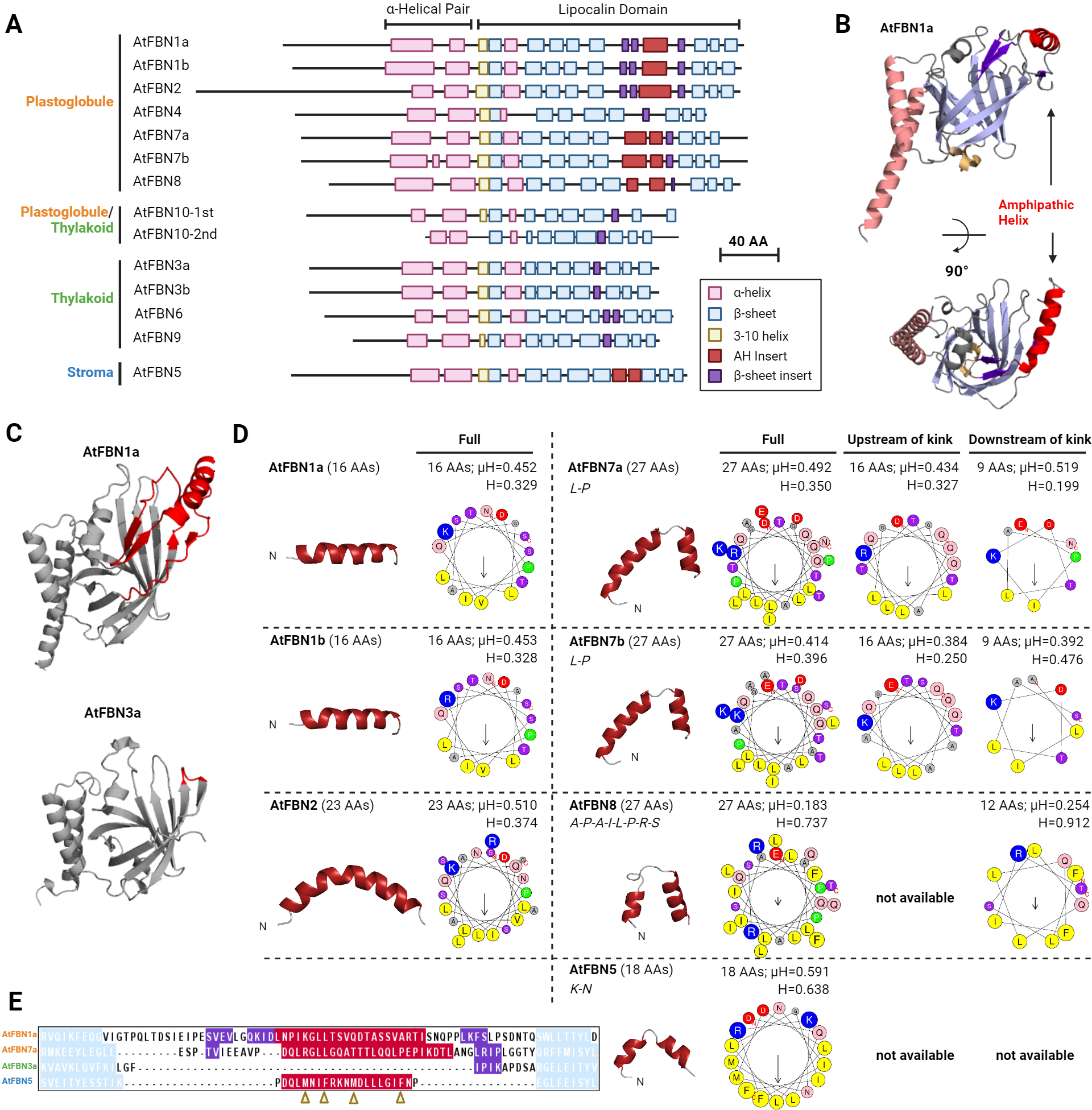
Predicted Structures of the *A. thaliana* Fibrillin Family. **A**, Schematic of secondary structures of all thirteen *A. thaliana* FBN proteins, as predicted by AlphaFold. The schematic is grouped by sub-plastidic localization and each sequence is drawn to scale and aligned based on the first β-sheet of the lipocalin domain which contains the absolutely conserved GxW motif. The 3-11 helix is presumed based on the helical structure of the AlphaFold model and the solved structures of other lipocalin domains. Note that AtFBN10 contains two repeated lipocalin domains which are illustrated on separate lines in the schematic; where the first sequence drops off, the sequence is then directly continued on the second sequence without un-captured gaps. **B**, AlphaFold-predicted structure of AtFBN1a. Secondary structural features are colored according to the schema used in panel A. **C**, Structural comparison of the plastoglobule-localized AtFBN1a and the thylakoid-localized AtFBN3a. Sequence highlighted in red indicates the 45-residue sequence surrounding the predicted AH in AtFBN1a and the corresponding sequence region in AtFBN3a which lacks any amphipathic helical insert. **D**, AlphaFold-predicted structures of amphipathic helical inserts from each FBN protein. All structures are presented at the same size scale and with N-terminal portion on the left. To the right, predicted amphipathic helical structure is presented for each HeliQuest-predicted insert sequence. Where relevant, predicted disordered sequence interruptions within the helical structure are included in the full-length analysis, and sub-sequences upstream or downstream of the interruption are additionally included below the protein name. The one-letter residue-code size is proportional to residue side chain volume. **E**, Sequence alignment of the amphipathic helical insert region from selected FBN proteins. The alignment highlights sequence differences, in particular the absence of large hydrophobic side chains from AtFBN1a and AtFBN7a, which are present in the non-plastoglobule-localized AtFBN5 AH and noted with gold arrowheads under the alignment.

The predicted AHs of the FBNs corresponded in some instances to the helical inserts (AtFBN1a, AtFBN1b, AtFBN7b) and in other instances with portions of the helical pair (AtFBN1b, AtFBN4). Because the presence of a helical insert closely coincided with plastoglobule localization, we considered how the helical inserts may drive interaction with plastoglobules. Analyzing the helical inserts of each of the seven FBNs, we found that they all held amphipathic character, with hydrophobic moments (µH) ranging from 0.414 to 0.591, except for AtFBN8 whose helical insert is interrupted by eight-residues which are predicted by AlphaFold to lack helical structure (**Fig. 1E**). Large hydrophobic residues (Trp, Phe, Tyr) were conspicuously absent from the hydrophobic faces of the helical inserts, except for AtFBN5 and AtFBN8 (**Fig. 1B, E**). This contrasts with AHs of cytosolic LD proteins that frequently contain large hydrophobic residues with prominent roles in LD localization (38, 40, 65, 66).

### Plastoglobule-localized FBN proteins require the AH insert for proper plastoglobule association

To investigate the role of the helical inserts in chloroplast localization, we selected two plastoglobule-localized FBNs, AtFBN1a and AtFBN7a, with a continuous and an interrupted helical structure, respectively. We developed a series of three deletions within the AtFBN1a sequence which removed, i) the downstream two-thirds of the helical insert and five downstream flanking residues (AtFBN1a^Δ17α^), ii) all of the insert plus ten upstream flanking residues (AtFBN7a^Δ30α^), and iii) all of the insert and fifteen flanking residues upstream and downstream (AtFBN7a^Δ45α^) (**Fig. 2A**). Similarly, a series of three deletions within the AtFBN7a sequence were developed, which removed, i) the upstream portion of the helical insert (AtFBN7a^Δ17α^), ii) the downstream portion of the helical insert (AtFBN7a^Δ10α^), or iii) all of the insert plus two flanking residues upstream and downstream (AtFBN7a^Δ27α^) (**Fig. 2A**). AlphaFold structure prediction of deletion variants, followed by structural alignments to the wild-type variant, indicated that the deletions would not impact the core structure of the proteins including the β-barrel and α-helical pair (**Fig. 2B**).

**Figure 2.**
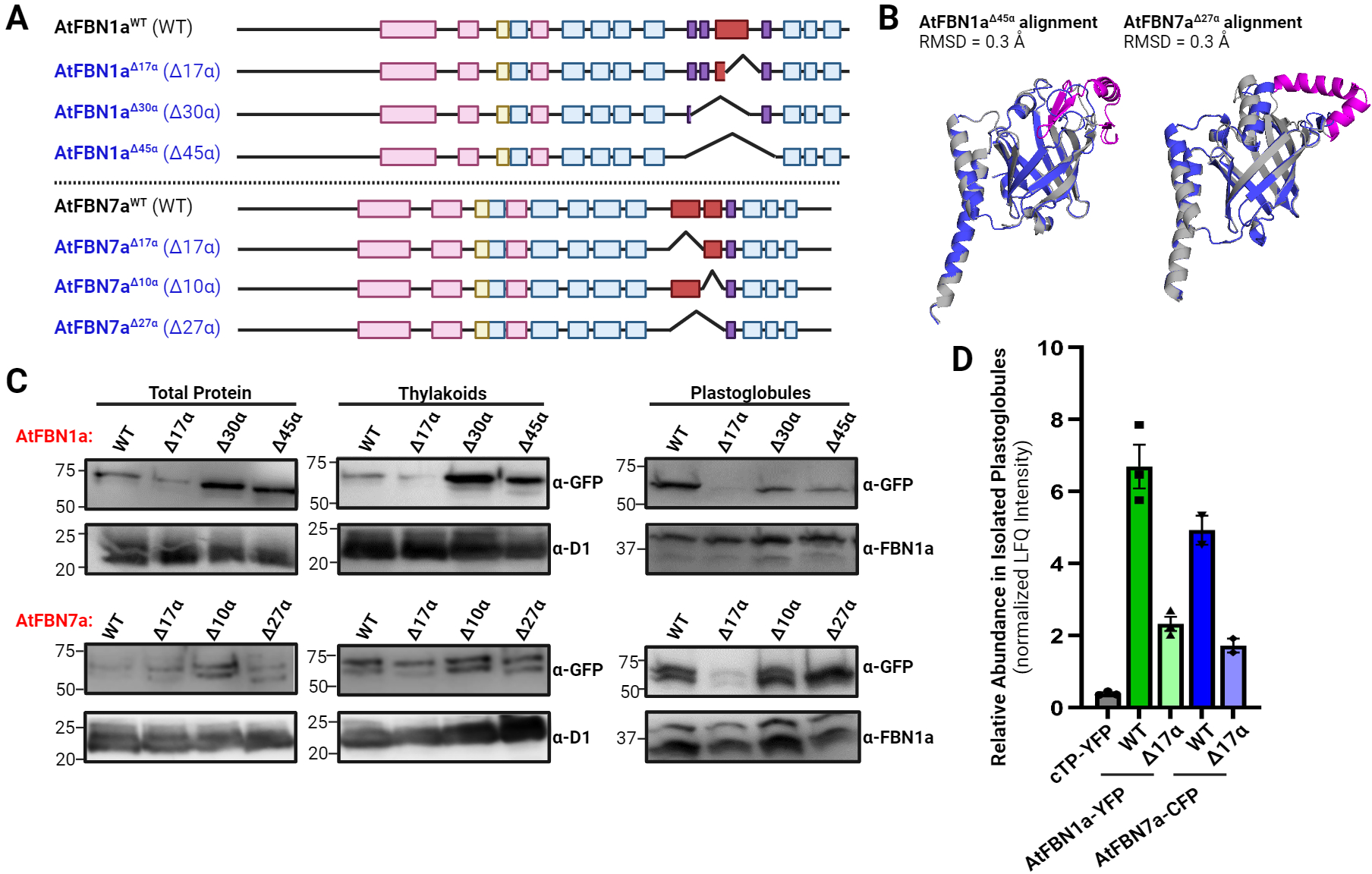
Deletion of the Predicted AHs in AtFBN1a and AtFBN7a Leads to Mis-localization Within Chloroplasts. A series of deletions were created in AtFBN1a and AtFBN7a transiently transformed into tobacco (N. benthamiana) and tracked by sub-chloroplast fractionation and immunoblotting. **A**, Schematics of the series of deletion mutants in the AtFBNs is presented along with amphipathic helical wheels of the predicted AHs of AtFBN1a and AtFBN7a. H, hydrophobicity; μH, hydrophobic moment. **B**, AlphaFold-predicted structural models of the Δ17α deletion variants were aligned with their corresponding wild-type structure. Alignments indicated that deletions are not expected to impact the overall sequence structure of the proteins, supported by circular dichroism data of heterologously expressed proteins (**Supp Fig. S4**). All-atom root mean square deviation (RMSD) was calculated with PyMol. **C**, *N. benthamiana* was transiently transformed with constructs expressing wild-type or deletion-variant sequences of AtFBN1a or AtFBN7a containing a C-terminal FP-tag. Two days after infiltration, thylakoid (unsonicated - thus plastoglobules have not been stripped off), and plastoglobule sub-compartments were isolated for immunoblotting and proteomics. Immunoblots using rabbit anti-GFP were used to monitor the AtFBN constructs. Rabbit anti-D1 and rabbit anti-FBN1a were used to monitor D1 (PsbA) and total FBN content, respectively, as loading controls. **D**, LC-MS/MS-based proteomic analysis using label free quantification (LFQ) of isolated plastoglobules.

Sequences were tagged with fluorescent protein (FP) and transiently transformed individually into tobacco plants (*Nicotiana benthamian*a), driven by the strong tobacco mosaic virus 35S constitutive promoter. Forty-eight hours after transformation, plastoglobule and thylakoid sub-compartments were isolated from leaf tissue and immunoblotted with anti-GFP antibody (**Fig. 2C**). As expected, the wild-type variants, AtFBN1a^WT^-YFP and AtFBN7a^WT^-CFP, were found strongly associated with the isolated plastoglobules. In contrast, full removal of the insert of AtFBN1a, as tested in AtFBN1a^Δ30α^-YFP and AtFBN1a^Δ45α^-YFP, resulted in a shift in localization to the thylakoid membrane. Partial deletion of the insert, in AtFBN1a^Δ17α^-YFP appeared to reduce the stability of the protein, as the shift to thylakoids was not seen as well as relatively low protein accumulation in total leaf tissue. A somewhat different picture emerged with AtFBN7a. Deletion of the upstream 17 residue AH, as tested in AtFBN7a^Δ17α^-CFP, severely reduced the plastoglobule association, with a corresponding shift to thylakoids (**Fig. 2C**). However, deletion of the downstream portion of the helical insert (AtFBN7a^Δ10α^-CFP) did not affect plastoglobule association and simultaneous deletion of both the upstream and downstream portions of the helical insert (AtFBN7a^Δ27α^-CFP) partially restored localization to the plastoglobule.

To validate our immunoblotting results, we used LC-MS/MS proteomics with label-free quantification to estimate the relative abundance of the wild-type and the Δ17α variant of AtFBN1a and AtFBN7a on isolated plastoglobules (**Table S2**). Consistent with the immunoblotting results, both Δ17α variants were reduced to about 1/3 the abundance of their respective wild-type variants (**Fig. 2D**). Nonetheless, levels of the deletion variants were significantly greater than the negative control, a stromal-localized YFP driven by the AtRbcS chloroplast transit peptide (cTP^AtRbcS^-YFP), indicating that appreciable amounts of AtFBN1a^Δ17α^-YFP and AtFBN7a^Δ17α^-CFP were able to associate with plastoglobules even with a partial deletion of their respective amphipathic helical insert.

We also investigated localization of the constructs by confocal microscopy. Punctate fluorescence within chloroplasts has previously been used to indicate plastoglobule localization of proteins, including FBNs (16, 18, 29). However, because punctate patterns can arise within chloroplasts due to reasons other than plastoglobule association, such as liquid-liquid phase separation in the stroma (67), we co-transformed each test construct tagged with FP along with a plastoglobule marker protein (AtFBN1a^WT^-YFP or AtFBN7a^WT^-CFP) to demonstrate *bona fide* plastoglobule localization through co-localized fluorescence. Unexpectedly, all co-transformations, including deletion variants, resulted in co-localized punctate patterns (**Fig. S4**). Punctate fluorescence was frequently observed on the edge of the chlorophyll auto-fluorescence, suggesting protein constructs were accumulating at the TOC-TIC import machinery. Confocal observations at later (72 hrs) or earlier time points (24 hrs) after transformation gave similar results to our 48 hr observations. We deemed the confocal microscopy analysis as inconclusive, possibly due to overwhelming demand on the chloroplast import apparatus when co-transforming with two plastid-targeted proteins. Based on the immunoblot and LC-MS/MS analyses of isolated plastoglobules and thylakoids, we conclude that the amphipathic helical inserts are necessary for effective association of FBN proteins with plastoglobules.

### MD simulations reveal differential interactions with thylakoid and plastoglobule surfaces

MD simulations were used to develop molecular models of AtFBN1a^WT^ interaction with membrane surfaces corresponding to plastoglobules or thylakoids. The plastoglobule surface was simulated using a tri-layer system comprising an 8 nm thick neutral lipid core sandwiched between two glycerolipid monolayers, as used in MD simulations with other LDs (**Fig. 3A**) (37, 68, 69). To mimic the plastoglobule in our simulations we utilized a galactolipid monolayer composition of equal parts monogalactosyl diacylglycerol (MGDG) and monogalactosyl monoacylglycerol (MGMG), derived from our lipidomic profiling of isolated plastoglobules (Devadasu & Lundquist - unpublished). The thylakoid was simulated with a bilayer membrane comprised of MGDG, digalactosyl diacylglycerol (DGDG), and sulfoquinovosyl diacylglycerol (SQDG) (**Fig. 3A**). Six unique simulation systems (replicas) were created with each membrane system using the AlphaFold structural model of AtFBN1a^WT^ (lacking the 68-residue predicted chloroplast transit peptide [-cTP]) rotated to face each side of a cube (**Fig. S5**), creating independent starting configurations to minimize bias. These starting configurations were run for 1.0 µs using NAMD 3.0a12 (70) and CHARMM36 force fields (71–73).

**Figure 3:**
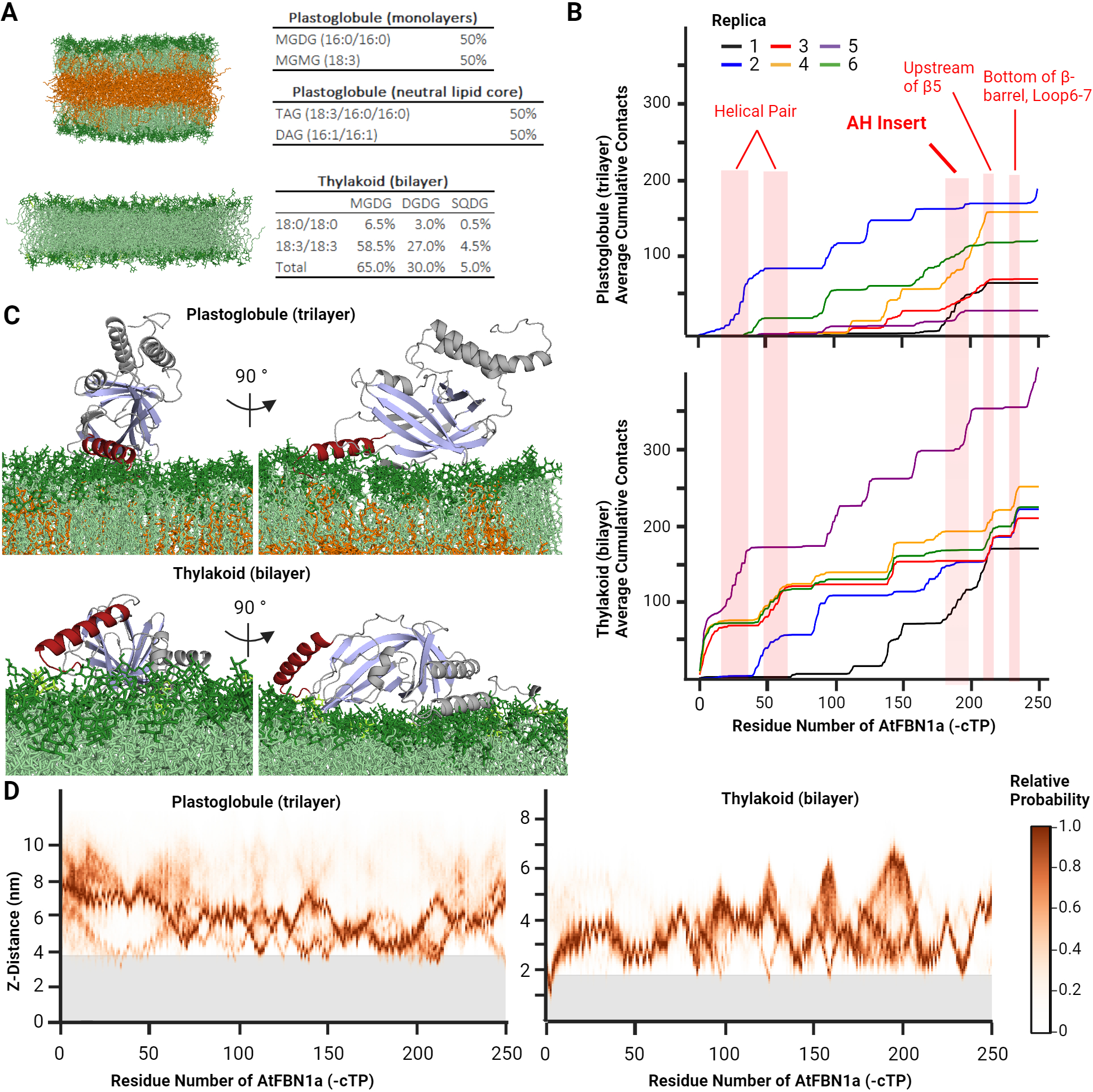
Molecular Dynamics Simulations of AtFBN1a Support an Amphipathic Helical Insert-Driven Mechanism of Plastoglobule Interaction. **A,** Illustration of the simulated membranes, a trilayer system mimicking the plastoglobule lipid droplet, and a bilayer mimicking the thylakoid membrane. Galactolipid headgroups are colored in dark green, sulfolipid headgroups in lime green, acyl tails in light green, and tri-and di-acylglycerol neutral lipids in orange. **B**, Cumulative contacts between protein and membrane, as measured by equation 1 in Experimental Procedures. The contacts between individual amino acids and the membrane are reported as a cumulative sum, to better visualize the overall effect of different protein regions on the binding interaction. The values are averaged between 500 and 1000 ns of trajectory for each replica. **C,** Snapshots of the simulations of Replica 2 (plastoglobule) and Replica 5 (thylakoid) at ca. 1.0 μs. Coloring of membrane lipids is as in **A**, and coloring of protein is as in Figures 1 and 2. **D**, Probable insertion depth is plotted as a function of residue. The 0 nm Z-distance is set as the midpoint of the membrane (gray). The top of the gray banner corresponds approximately to the top of the lipid headgroups.

Membrane contact points along the protein sequence were identified and mapped (**Fig. 3B**). Highly distinct binding poses emerged in the presence of simulated plastoglobule and simulated thylakoid membranes. In the presence of the plastoglobule trilayer, membrane contacts primarily arose along the AH insert and the downstream 15 residues (residues 195 to 209 of the full-length sequence); a binding pose which emerged in five of the six replicas (**Fig. 3B and C**). No other regions consistently contacted the plastoglobule across multiple replicas. In contrast, in the presence of the thylakoid bilayer, contacts were identified at the N-terminus of the mature protein sequence (residues 1 to 10 of the full-length sequence) which contained several bulky hydrophobic side chains (3V, 4F, and 10V), the downstream α-helix of the helical pair, upstream of β-strand E, and the βF-βG loop, while the AH insert only contacted the thylakoid surface in two of the six replicas (**Fig. 3B, C, & D**). Notably, binding depth of the protein did not substantially penetrate below the polar headgroups in the plastoglobule or thylakoid simulations, with the exception of the N-terminus on the thylakoid bilayer (**Fig. 3D**). This suggests that either the 1.0 µs trajectories were not long enough for residues to penetrate more deeply into our plastoglobule mimic, or that plastoglobule curvature is required for residue intercalation. Further investigation with substantially larger plastoglobule LDs would facilitate discrimination between these two hypotheses. Nonetheless, our MD simulation results support a central role for the AH insert in plastoglobule interaction via the unique plastoglobule lipid composition and structure.

### Accumulation of exogenous FBN selectively disrupts the plastoglobule proteome

Localization of certain proteins on cytosolic LDs is sensitive to protein crowding, in which such proteins are outcompeted by stronger-binding proteins (33, 38, 74). Furthermore, plastoglobule localization of certain proteins may be dependent on protein-protein mediated recruitment (3, 4, 26, 27). To investigate whether AtFBN1a may influence the plastoglobule proteome through protein crowding or protein recruitment we characterized the quantitative proteome of plastoglobules using the LC-MS/MS analyses of isolated plastoglobule from *N. benthamiana* transient transformations described above. Isolated plastoglobules from the cTP^AtRbcS^-YFP-accumulating tissue represent the negative control and provide a baseline proteome of *N. benthamiana* plastoglobules which has not previously been reported in the literature. Our objective was not to identify novel plastoglobule-localized proteins, but to identify and quantify the expected plastoglobule-localized proteins of *N. benthamiana* and how they are changed in abundance. Therefore, we focused our identification and quantification on orthologs of the *A. thaliana* plastoglobule proteins identified from either un-stressed or light-stressed conditions (7, 15). Most of the orthologs were identified in the isolated *N. benthamiana* isolated plastoglobules of the negative control, comprising 20 distinct proteins or protein groups (hereafter, proteins) (**Table 1; Table S2**). An additional four proteins were identified in either the AtFBN1a^WT^-YFP or AtFBN1a^Δ17α^-YFP plastoglobules. Protein orthologs not identified in any *N. benthamiana* plastoglobule isolation were limited to Esterase/Lipase/Thioesterase 4, Allene Oxide Synthase, Aldo/Keto Reductase, and several proteins with unannotated function (7, 15). These proteins are amongst the lowest abundant proteins of *A. thaliana* plastoglobules and, in some cases only localized to *A. thaliana* plastoglobules under light-stress conditions. This indicates that they may be present on *N. benthamiana* plastoglobules, albeit only under stressed conditions or below the limit of detection of our experimental setup.

Broadly speaking, the baseline plastoglobule proteome, represented by the cTP^AtRbcS^-YFP accumulating samples, is similar to that of unstressed *A. thaliana*, characterized by several abundant FBN homologs, particularly FBN4, and a number of lower abundant enzymes including several homologs of the ABC1K family (**Fig. 4A**; **Table 1**). However, it is notable that the proportion of the *N. benthamiana* plastoglobule proteome dedicated to FBNs was substantially greater than that seen in *A. thaliana* (68% in *N. benthamiana* vs. 44% in *A. thaliana*). Additionally, Plastoglobular Protein of 18 kD (PG18), SOUL Heme-Binding Protein 3 (HBP3) and Carotenoid Cleavage Dioxygenase 4 (CCD4) were considerably more abundant in *A. thaliana* plastoglobules (1.0%, 1.0%, and 0.6% in *N. benthamiana* vs. 5.2%, 9.6%, and 4.9% in *A. thaliana*, respectively).

**Figure 4:**
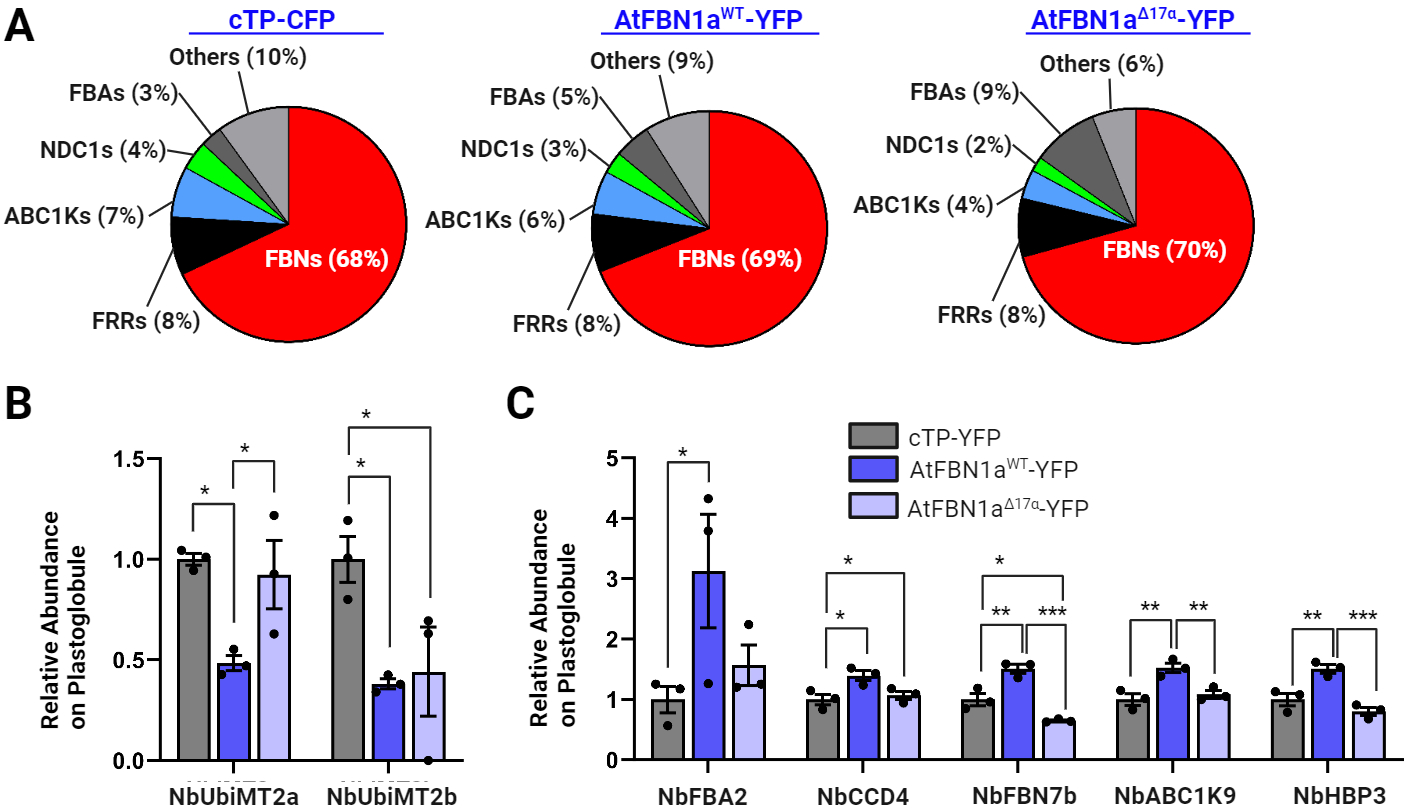
Impacts of AtFBN Protein Variants on the *N. benthamiana* Plastoglobule Proteome. The *N. benthamiana* plastoglobule proteome was assessed using LC-MS/MS of plastoglobules isolated from leaf tissue expressing either the wild-type or Δ17α variant of AtFBN1a, tagged C-terminally with YFP (AtFBN1a^WT^-YFP and AtFBN1a^Δ17α^-YFP, respectively), or YFP alone with the RbcS chloroplast transit peptide (cTP) to ensure chloroplast import (cTP-YFP). **A,** Twenty-eight proteins were classified as likely plastoglobular based on their enrichment in isolated plastoglobules and homology to members of the *A. thaliana* plastoglobule proteome. Their relative abundance is presented by protein family grouping. **B, C,** Bar graphs indicate the relative abundance of selected proteins on isolated plastoglobules, normalized to 1 when expressing the cTP-YFP variant. n = 3, mean ± 1 s.e.m. Statistically significant differences determined by a standard one-way ANOVA are indicated, * <0.033, ** <0.002, *** <0.001.

We found that over half of the *N. benthamiana* endogenous proteome changed in abundance when either AtFBN1a^WT^-YFP or AtFBN1a^Δ17α^-YFP were localized on plastoglobules (**Table 1**). We considered proteins as putatively recruited to plastoglobules by AtFBN1a if they were increased in abundance in AtFBN1a^WT^-YFP samples, but increased to a lesser extent (or not at all) in AtFBN1a^Δ17α^-YFP samples. Conversely, we considered proteins as putatively outcompeted on the plastoglobule by AtFBN1a if they were decreased in abundance in AtFBN1a^WT^-YFP samples, but decreased to a lesser extent (or not at all) in AtFBN1a^Δ17α^-YFP samples. According to these guidelines, we identified two proteins that were putatively outcompeted by AtFBN1a: i) UbiE-like Methyl-transferase 2a (NbUbiMT2a), and ii) UbiE-like Methyl-transferase 2b (NbUbiMT2b) (**Fig. 4B**, **Table 1**). Likewise, we identified five proteins that were putatively recruited to the plastoglobule by AtFBN1a: i) Fructose Bis-phosphate Aldolase 2 (NbFBA2), ii) NbCCD4, iii) NbFBN7a, iv) NbABC1K9, and v) NbHBP3 (**Fig. 4C**, **Table 1**). The strongest recruitment appeared to be for NbFBA2 which increased in abundance by 3-fold specifically in the presence of AtFBN1a^WT^-YFP. Notably, our results provided some support for possible recruitment of NbABC1K3 as well, although the low abundance of the protein meant the difference between AtFBN1a^WT^-YFP and cTP^AtRbcS^-YFP was not statistically significant. Curiously, our results indicated that NbPG18 was lost from plastoglobules when expressing AtFBN1a^Δ17α^-YFP. It is not clear what relationship the deletion variant of AtFBN1a may have on NbPG18 localization. Collectively, our proteomic investigation uncovers evidence that ectopic expression of AtFBN1a alters the plastoglobule proteome, at least in part *via* protein recruitment and competition.

### The AH of AtFBN1a is ineffective at re-directing thylakoid-or stromal-associated FBN protein to the plastoglobule

To test the sufficiency of the AtFBN1a AH for plastoglobule localization, we introduced the AH into the loop regions between β-strands E and F of AtFBN3a and AtFBN5, localized to the thylakoid and the stroma, respectively. To ensure adequate flexibility of the inserted AH and account for possible roles in plastoglobule association, we included upstream and downstream flanking sequence for a total of 45 residues of AtFBN1a (residues 232 – 276 of the full-length sequence) (**Fig. 5A**). As for the deletion variants, insertion variants were tagged C-terminally with FP and tested for sub-chloroplast localization using immunoblotting of isolated sub-compartments. As expected, AtFBN3a^WT^-CFP associated predominantly with isolated thylakoid (**Fig. 5C**). However, the introduction of the insert sequence re-directed AtFBN3a^+45α^-CFP to the stroma and envelope, with no detectable accumulation on the plastoglobule of either the WT or +45α variants. Similarly, introduction of the AH to AtFBN5 did not lead to any detectable accumulation with isolated plastoglobules (**Fig. 5C**). We also attempted to target constructs in which the 45-residue sequence was placed at the C-terminus of the AtFBN3a and AtFBN5-A proteins (upstream of the C-terminal FP tag), however protein accumulation *in planta* was very low and insufficient to judge sub-cellular localization. We conclude that introduction of the 45-residue sequence containing the AtFBN1a AH is not sufficient to redirect these two FBN proteins to the plastoglobule, and that its position in the protein sequence can affect protein stability.

**Figure 5:**
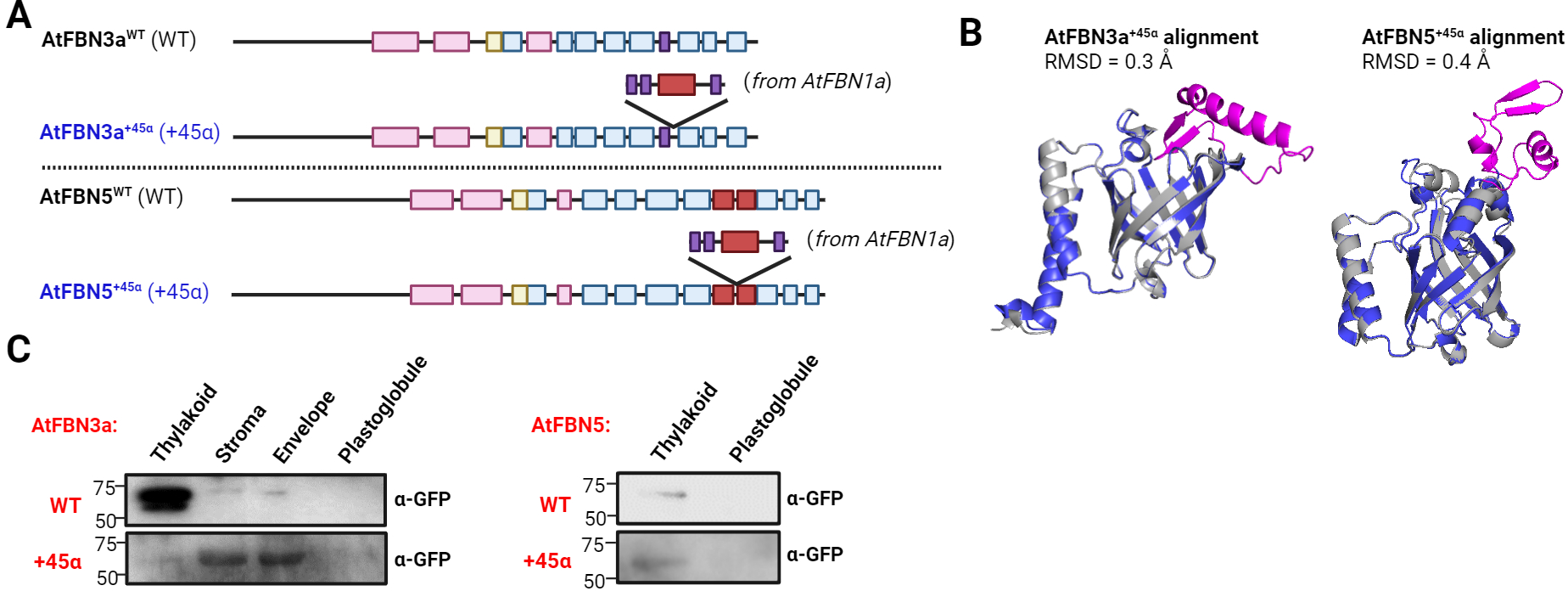
Inclusion of an AH in Thylakoid or Stromal FBNs Does Not Redirect Localization to the Plastoglobule. **A,** Sequence variants of AtFBN3a and AtFBN5 were created by inserting the 45 residues sequence of the AtFBN1a AH and flanking sequence between β-strands E and F. **B**, AlphaFold-predicted structural models of the +45α insertion variants were aligned with their corresponding wild-type structure. Alignments indicated that insertions are not expected to impact the overall sequence structure of the proteins (**Supp Fig. S4**). All-atom root mean square deviation (RMSD) was calculated with PyMol. **C**, *N. benthamiana* was transiently transformed with constructs expressing wild-type or the insertion-variant sequences, AtFBN3a^+45α^ or AtFBN5^+45α^, containing a C-terminal CFP-tag on the AtFBN3a variants and YFP-tag on the AtFBN5 variants. Two days after infiltration, thylakoid and plastoglobule sub-compartments were isolated for immunoblotting. Immunoblots using rabbit anti-GFP were used to monitor the presence of AtFBN constructs.

### FBN proteins display lipid-specific binding that is affected by loss of the AH insert

The lipocalin domain, characteristic of the FBN family, is capable of binding a wide array of small hydrophobic compounds, particularly free fatty acids (19, 20, 75, 76). While the lipocalin sequence of FBNs does not share all characteristic motifs of the canonical lipocalin domain (77), for example the large loop L1 is absent in FBNs, the eight-stranded β-barrel structure is reproduced in the AlphaFold and RoseTTAFold structural models and the GxW motif is conserved at the bottom of the barrel at the beginning of the βA strand. To investigate the lipid binding abilities of AtFBN1a and AtFBN7a, and the impact of the Δ17α sequence variants lacking the core of the predicted AHs, we employed *in vitro* assays using heterologously expressed protein with the P-96 polarity-sensitive fluorescent probe (78). P-96 fluoresces strongly in a hydrophobic environment, in our case provided by binding to the AtFBN protein. Competitive binding by an interacting lipid species will dislodge the P-96, reducing the fluorescence signal and giving a quantitative readout of lipid-binding efficacy. Based on the binding specificities of other lipocalin domains and the composition of the plastoglobule, we assayed a series of free fatty acids (FAs), including saturated and unsaturated acyl chains of varying length, as well as phosphatidic acid and diacylglycerol species.

The secondary structure of heterologous expressed proteins was validating using circular dichroism (CD) spectroscopy. Experimentally calculated secondary structure of each protein closely aligned with prediction based on the AlphaFold models (**Table 3**). Further, the deletion variants showed the expected reduction in proportion of helical structure. Because CD spectra were collected in aqueous solution in the absence of liposome, we conclude that the AHs adopt a stable helical structure even in the absence of membrane. AtFBN1a^WT^ showed a clear preference for unsaturated FAs, with especially effective binding of C18:1 (**Fig. 6**). Notably, binding efficacy of lipids was impaired in the absence of the AH. AtFBN7a^WT^ also displayed a clear preference for C18:1 although binding efficacy was less than that of AtFBN1a^WT^. Strikingly, the loss of the AH from AtFBN7a promoted binding of lipids, the opposite effect of AH deletion in AtFBN1a. As an orthogonal approach we employed lipid overlay assays to test lipid binding specificity (**Fig. S6**). AtFBN7a^Δ17α^, but not AtFBN7a^WT^, interacted specifically with unsaturated FAs, consistent with the results from the P-96 fluorescence assays. However, the lipid overlay assays did not detect interaction of AtFBN1a variants with any free FAs; the more artificial manner in which lipid species are presented in the lipid overlays may impact AtFBN1a’s ability to stably bind lipid. Collectively, our results indicate that both proteins are capable of binding lipid species, particularly C18:1, and that the AH influences lipid binding in opposing manners in AtFBN7a and AtFBN1a.

**Figure 6.**
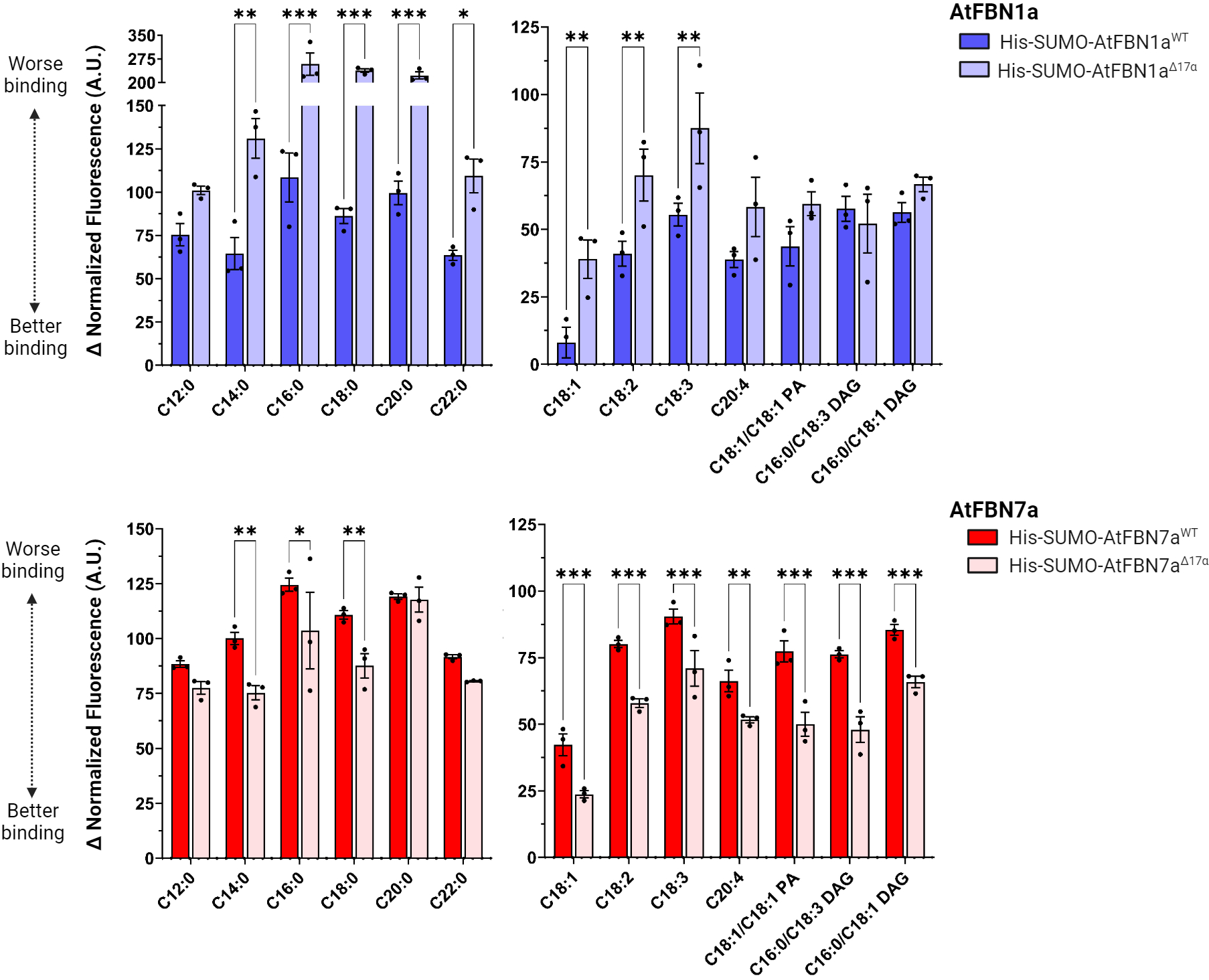
Lipid-binding of plastoglobule-localized AtFBN1a and AtFBN7a proteins. Lipid-binding was assayed using an in vitro assay setup with the polarity-sensitive hydrophobic fluorophore, 1-Pyrenedodecanoic acid (P-96). Various free fatty acids, phosphatidic acid (PA) and diacylglycerol (DAG) species were assayed with wild-type and Δ17α sequence variants of AtFBN1a and AtFBN7a. Competitive binding that displaced P-96 was represented by a reduction in fluorescence.

**Table 3.**
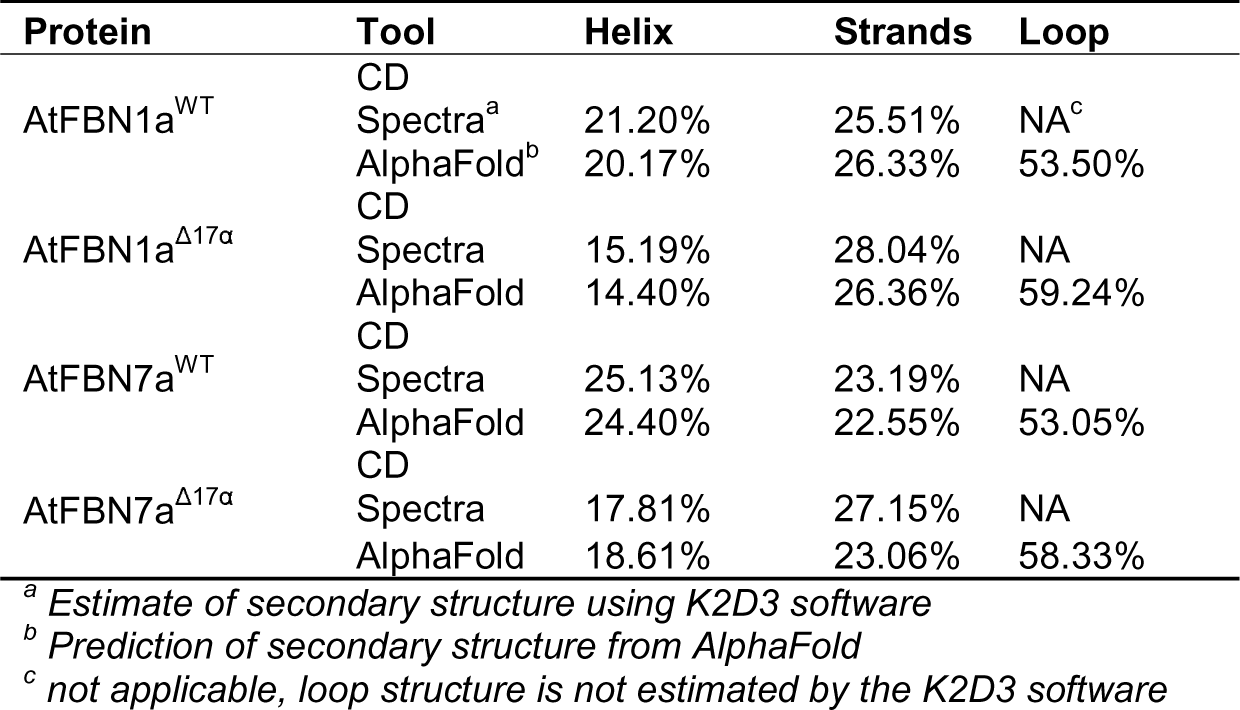
Estimation of Secondary Structure of Heterologously.

## DISCUSSION

The mechanisms by which specific proteins associate with plastoglobule LDs remain obscure. In light of the plastoglobule’s monolayer structure enclosing a neutral lipid core, transmembrane helices cannot be accommodated; rather, associated proteins should presumably interact in a monotopic fashion, either peripherally or *via* hairpin loop. But how can proteins distinguish between plastoglobules and the other membrane systems of the chloroplast: the extensive internal thylakoid membranes and the encompassing inner envelope membrane? In the present work, we have demonstrated the significance of a conserved AH in driving plastoglobule localization of two members of the FBN family.

Our discovery that the AH insert of AtFBN1a is responsible for direct interaction with plastoglobules is consistent with our previous study in which we used protease shaving of isolated plastoglobules to map exposed and protected regions of the protein (79). This work suggested a tryptic cleavage site between the α-helical pair and the β-barrel which, together with phosphosites on either side of the α-helical pair identified from publicly available phosphoproteomic datasets, led us to propose that the α-helical pair is positioned distal to the plastoglobule surface and the AH insert in contact with the plastoglobule. In work by Vidi, et al., confocal microscopy of a series of truncated variants of AtFBN7a indicated nearly the entire protein sequence was necessary for plastoglobule association (residues 1-290 of the full length AtFBN7a) (29). Shorter variants that still retained the AH insert (sequences 224-250 of the full-length AtFBN7a) appeared to aggregate within the chloroplast and failed to associate with plastoglobules. From the AlphaFold-modeled structure we can see that those variants would have disrupted the β-barrel, likely leading to the observed aggregation. Thus, the truncation study is consistent with our reported significance of the AH inserts and indicates that a targeted deletion of the AH and flanking sequence is necessary to abolish plastoglobule localization while maintaining overall protein structure and preventing aggregation.

There is precedent for the use of AHs to target proteins to cytosolic LDs (33, 38, 50, 51, 80). However, there are a number of important distinctions between plastoglobules and cytosolic LDs that may confound the direct exchange of targeting principles (1). First, the polar lipid composition of the plastoglobule is surrounded by neutral galactolipids (2, 81–83), in contrast to the phospholipid-rich nature of cytosolic LDs which includes negatively charged phosphatidylglycerol and phosphatidylcholine (84, 85). Interactions between the negative charge of the cytosolic LD surface and the positively charged residues located at the hydrophobic-hydrophilic interface of AHs can be important for AH interaction with cytosolic LDs (42). However, such interactions would seemingly not be significant on the plastoglobule surface with their only limited negative charge from low levels of SQDG. Second, the composition of the neutral lipid core is highly distinct between cytosolic LDs and plastoglobules. Whereas cytosolic LDs are dominated by TAG and/or sterol ester compounds (1, 44, 45), plastoglobules hold only low levels of TAG and lack sterols, instead being enriched in phytyl esters and prenyl-lipids (10, 13, 86). Specific interactions between protein and TAG can promote protein association on cytosolic LDs (66, 87); different protein-lipid interactions may drive targeting to plastoglobules. Third, the smaller size of the plastoglobules (often as little as 70 nm in diameter) compared to that of cytosolic LDs (sometimes exceeding a thousand nm in diameter) introduces differing degrees of curvature to LD surfaces which can affect the magnitude and nature of the lipid packing defects. Thus, the underlying neutral lipids may be particularly susceptible to emergence on the plastoglobule surface.

Consistent with the multiple distinctions between plastoglobules and cytosolic LDs, bulky hydrophobic side-chains are conspicuously absent from AHs of the plastoglobule-targeted FBNs. This contrasts with AHs of cytosolic LD proteins in which bulky hydrophobic side chains are typically present and are important for proper association (34, 38–41). On the other hand, FBN AHs share an enrichment of polar side-chains, particularly near the hydrophobic-hydrophilic interface (**Fig. 1**), as seen with ALPS-like AHs found in curvature-sensing proteins of cytosolic LDs (36, 42). This indicates that a similar principle may be used for curvature sensing between cytosolic LDs and plastoglobules. Collectively, the sequences of the FBN AHs and distinct size and composition of plastoglobules underline the likelihood that biophysical principles differ to some extent between plastoglobules and cytosolic LD protein association.

Our use of MD simulations support direct interaction of the AH inserts with the plastoglobule, and the importance of lipid packing defects on the membrane surface to foster this interaction. There are caveats to the conclusions which can be drawn from these simulations. As shown in **Fig. 3B**, contact profiles are not perfectly consistent, indicating that 1 µs is not enough to fully sample rotational degrees of freedom for protein-membrane contacts. Longer simulation times may be necessary to establish stabilizing interactions between protein and membrane including putative specific polar and hydrophobic interactions. This would be valuable in understanding the molecular details of AH interaction on the plastoglobule surface by establishing the roles of specific residues, hypothesizing effects from the introduction of bulky hydrophobic side chains, and estimating the strength of hydrophobic interactions in AH insertion.

The position of the AH insert near the opening of the β-barrel may hold implications for protein function and lipid binding. Our lipid-binding assays indicate that the presence or absence of the AH insert effects lipid binding strength, albeit in opposing directions among the two tested proteins. This puzzling result may lie in the different AlphaFold-predicted structures of the AH inserts of AtFBN1a, which has a single contiguous AH, and AtFBN7a, which has two non-contiguous AHs. The tested deletion constructs remove all of the AtFBN1a AH, but only the upstream portion of the AtFBN7a AH. It may be that the AH inserts represent lids covering the opening of the hydrophobic cavity of the β-barrel that prevent lipid binding until interaction with membrane causes the AH insert to move. This would present the AH insert as a sort of gate for lipid binding that is only opened on the proper membrane surface. The lipid binding capacity of the FBNs also raises the question of what purpose specific lipid binding may serve, including possible involvement in plastoglobule targeting. Arguing against a role in plastoglobule targeting, we show that Δ17α variants of AtFBN1a and AtFBN7a hold opposing effects on lipid binding strength (**Fig. 6**), but similarly impair plastoglobule association (**Fig. 2**), observations inconsistent with a role for lipid binding in plastoglobule targeting. More plausibly, FBNs may function as lipid transporters, shuttling specific lipid species between membrane systems, as has been proposed previously (88).

## EXPERIMENTAL PROCEDURES

### Transient expression in *Nicotiana benthamiana*

The full length FBN proteins were cloned into either pEARLEYGate101 or pEARLEYGate102 vectors to transiently express FBN protein with fluorescent tagged proteins. The chloroplast transit peptide of AtRbcL and the full length and truncated AtFBN1a variants, including their chloroplast transit peptides, were cloned as coding sequences into pEARLEYGate101 providing a C-terminal EYFP tag driven by 35S promoter. AtFBN7a, AtFBN3a and AtFBN5, and their mutated variants, including the chloroplast transit peptide, were cloned as coding sequences into pEARLEYGate102 vector providing a C-terminal ECFP tag and driven by 35S promoter. Constructs were transformed into agrobacterium AGL1 cells. Different combinations of constructs were infiltrated into 4-5-week-old *N. benthamiana* plants grown on Arabidopsis Mix soil at 25^0^ C, 100 μmol m^2^ s^-1^ and 16/8 hrs light/dark photoperiod. Infiltration of whole *N. benthamiana* plants was performed by applying vacuum as described in Bibik et al., 2022 (89). Chloroplast sub-compartment isolation or confocal experiments were performed 48 hrs after infiltration.

### Sub-cellular fractionation and immunoblotting

From *N. benthamiana* plants expressing AtFBN1a, AtFBN5-A, and AtFBN7a variants, plastoglobules and thylakoid membranes were extracted according to Shivaiah et al., 2022 (90). Briefly, plastoglobules were separated using 0.2 M sucrose in Medium R solution (50 mM HEPES-KOH pH 8.0 and 5 mM MgCl2 with phosphatase and protease inhibitors) after sonicating thylakoid fractions. The floating PGs were further purified using a sucrose gradient ranging from 0.7 M to 0 mM Sucrose. Purified PGs were normalized by measuring OD_700_.

From *N. benthamiana* plants expressing AtFBN3a variants, plastoglobules, thylakoids, envelope and stroma fractions were isolated according to Espinoza-Corral et al., 2021 (7). Briefly, infiltrated tobacco leaves were ground with an isolation buffer composed of 13 mM Tris-HCl, 20 mM MOPS at pH 7.6, 3 mM MgCl_2_, 0.1% [w/v] BSA, 5 mM ascorbic acid, 5 mM reduced cysteine, 330 mM sorbitol, and a phosphatase inhibitor cocktail containing 50 mM NaF, 25 mM β-glycerophosphate, 1 mM Na-orthovanadate, and 10 mM Na-pyrophosphate. This mixture was then passed through gauze filter. Chloroplasts were separated by centrifugation at 1500 g for 5 minutes at 4^0^C, and subsequently washed twice with a washing buffer consisting of 50 mM HEPES at pH 7.6, 5 mM ascorbic acid, 5 mM reduced cysteine, 330 mM sorbitol, and a cocktail of phosphatase inhibitors.

The washed chloroplasts were re-suspended in an osmotic stress buffer containing 10 mM Tricine at pH 7.9, 1 mM EDTA, 0.6 M sucrose, and a mixture of phosphatase and protease inhibitors, including 74 μM antipain, 130 μM bestatin, 16.5 μM chymostatin, 56 μM E64, 2.3 μM leupeptin, 37 μM phosphoramidon, 209 μM AEBSF, 0.5 μM aprotinin, 50 mM NaF, 25 mM β-glycerophosphate, 1 mM Na-orthovanadate, and 10 mM Na-pyrophosphate. This chloroplast suspension was then incubated in darkness on ice for 30 minutes. After incubation, the broken chloroplasts were separated by centrifugation at 100,000 g for 1 hour at 4^0^C, resulting in the separation of the stroma from the membranes. The membranes were subsequently resuspended in 48% (w/v) sucrose in HE buffer (composed of 50 mM HEPES at pH 7.9, 2 mM EDTA, and a cocktail of protease and phosphatase inhibitors) and subjected to sonication with four pulses of 10 seconds each at 100% amplitude. The sonicated chloroplast membranes were overlaid with 5% (w/v) sucrose in HE buffer and then centrifuged at 150,000 g for 1.5 hours at 4^0^C. The plastoglobules were collected from top of the surface of the sucrose gradient, where they formed a yellowish band. Meanwhile, thylakoids and envelopes were collected form green pellet and light brown/white band at the interface of sucrose gradient, respectively.

For all sub-chloroplast isolations, proteins were quantified using a Pierce BCA Protein Assay Kit (Thermo Scientific) and an equal amount of protein was added to Laemmli sample buffer before boiling for 10 min. Samples were run on a 12% sodium dodecyl sulfate (SDS) gel and transferred to a nitrocellulose membrane (Amersham). Membranes were incubated with appropriate primary antibodies and then visualized using enhanced chemiluminescence (GE Healthcare).

### Confocal microscopy

Confocal experiments were performed on infiltrated *N. benthamiana* leaf tissue using a Nikon A1R-TIRF-STORM confocal microscope in the Center for Advanced Microscopy core facility on MSU campus. YFP fluorescence was monitored with an excitation wavelength of 514 nm and a bandpass of 530-600 nm emission filter, while CFP fluorescence was monitored at 485/35 (467-502 nm Band pass) and chlorophyll auto fluorescence was monitored at 660 mm long pass filter. Images were captured with, and analyzed using, Nikon NIS-Elements software.

### Protein structure modeling and molecular dynamics simulations

Alphafold was used to generate a protein structure from the amino acid sequence of AtFBN1a (56). This protein model was placed above a membrane model (in which each leaflet contained 400 lipids) in six different orientations and simulated using NAMD 3.0a12 to capture binding interactions (70). The bound states were analyzed to quantify the interactions between AtFBN1a and the membrane using VMD (91), highlighting the amino acids most responsible for its interaction.

Each of the six unique simulation systems were solvated in water using Python scripts built for the system. After solvation the system contained approximately 370,000 atoms. The dimensions of the simulation box are 158.8 Å by 158.8 Å by 144.4 Å. The simulation box was sized to permit realistic protein dynamics unconstrained by periodic boundary effects while reducing simulation cost. The system was briefly minimized and equilibrated using NAMD 2.14 to eliminate bad contacts (70). The simulations were performed using NAMD 3.0a12 (70) using explicit solvent and CHARMM36 carbohydrate, lipid, and protein force fields (71–73). In keeping with CHARMM standards, a 12 Å cutoff was used, together with the TIP3 water model (92). Long range electrostatics were handled using the particle mesh Ewald method with 1.2 Å grid spacing (93, 94). The system was simulated in an NPT ensemble using a Langevin thermostat and barostat (95). To enable 2 fs time-steps, we used the SETTLE algorithm to fix hydrogen bond lengths (96). The end simulation time for each individual replica was 1.5 µs. The simulations were carried out on NCSA’s Delta computing cluster.

Analysis was conducted using Python VMD scripts built for the system to leverage NumPy (97) for efficient numerics and Matplotlib (98) for plotting. Most of the analysis was conducted by analyzing intermolecular contacts present in the simulation, which measure the number of times protein residues approached the membrane within a certain cutoff distance. Depending on the cutoff distance selected, contact counts can change significantly. For our analyses, we use a weighted contact definition that allows for fractional contacts at distances near the selected cutoff. The exponentially weighted native contact definition from Shienerman and Brooks (99) has been used in prior studies to assess membrane-protein contacts (100–102). We define contact (*C*) as:

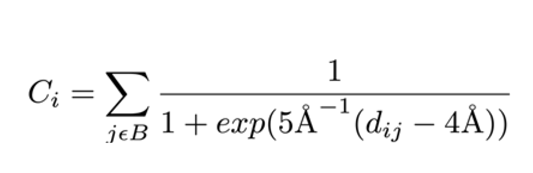

### LC-MS/MS proteomics and data analysis of plastoglobules

Isolated plastoglobules were prepared and analyzed by LC-MS/MS proteomics essentially as in Espinoza-Corral & Lundquist, 2021 (5). Lyophilized samples in triplicate (three biological replicates) were resuspended in 50 μl of 125 mM Tris, pH 6.8, 4% SDS, 20% glycerol, heated to 60 LJC for 5 min and electrophoresed into a 12% Tris-HCl Criterion gel from BioRad at 50 V for 20 min or until the dye front moved just below the bottom of the sample wells. Electrophoresis was stopped and the gel stained with Coomassie Blue protein stain until bands appeared. Protein bands were excised from the gel and digested according to Shevchenko et al. (103) with modifications. Briefly, gel bands were dehydrated using 100% acetonitrile and incubated with 10 mM dithiothreitol in 100 mM ammonium bicarbonate, pH 8, at 56 LJC for 45 min, dehydrated again, and incubated in the dark with 50 mM iodoacetamide in 100 mM ammonium bicarbonate for 20 min. Gel bands were then washed with ammonium bicarbonate and dehydrated again. Sequencing grade modified trypsin was prepared to 0.01 μg/μl in 50 mM ammonium bicarbonate and 100 μl of this was added to each gel band so that the gel was completely submerged. Bands were then incubated at 37 LJC overnight. Peptides were extracted from the gel by water bath sonication in a solution of 60% Acetonitrile (ACN)/1% Trifluoroacetic acid (TFA) and vacuum dried to 2 μl. Injections of 5 μl were automatically made using a Thermo (www.thermo.com) EASYnLC 1200 onto a Thermo Acclaim PepMap RSLC 0.1 mm × 20 mm C18 trapping column and washed for 5 min with Buffer A. Bound peptides were then eluted over 65 min onto a Thermo Acclaim PepMap RSLC 0.075 mm× 500 mm resolving column with a gradient of 8% B to 40% B in 54 min, ramping to 90% B at 55 min and held at 90% B for the duration of the run (Buffer A= 99.9% Water/0.1% Formic Acid, Buffer B = 80% Acetonitrile/0.1% Formic Acid/19.9% Water) at a constant flow rate of 300 nl/min. Column temperature was maintained at a constant temperature of 50 LJC using and integrated column oven (PRSO-V2, Sonation GmbH). Eluted peptides were sprayed into a ThermoScientific Q-Exactive HF-X mass spectrometer (www.thermo.com) using a FlexSpray spray ion source. Survey scans were taken in the Orbitrap (60,000 resolution, determined at m/z 200), and the top 15 ions in each survey scan are then subjected to automatic HCD with fragment spectra acquired at 7500 resolution.

The resulting MS/MS spectra were converted to peak lists using MaxQuant v2.4.5.0 and searched against a protein database using the Andromeda search algorithm, a part of the MaxQuant environment (104, 105). LC-MS/MS.raw files from samples expressing each protein variant were searched separately using a protein database containing all *N. tabacum* protein sequences available from UniProt (https://www.uniprot.org/uniprotkb?query=nicotiana+tabacum; 82358 proteins) with common laboratory contaminants and the respective AtFBN protein sequence appended. Oxidation of methionine, deamidation of asparagine, and N-terminal and lysine acetylation were set as variable modifications; carbamidomethylation was set as a fixed modification. Digestion mode was trypsin/P with a maximum of two missed cleavages. MS/MS tolerance of the first search was 20 ppm, and the main search was 4.5 ppm, with individualized peptide mass tolerance selected. FDR at peptide spectrum match and protein levels was set as 0.01, estimated using the reverse decoy method (106). LFQ intensity was determined as the total sum of the peak of Unique and Razor peptides with Fast LFQ selected. The best gene model for each protein group was identified as that with the highest number of Unique + Razor Peptide counts when compiled across all experiments. In the case of a tie, the lowest gene model number was selected, regardless of whether the protein group contained gene models from one or more gene loci.

The mass spectrometry proteomics data have been deposited to the ProteomeXchange Consortium via the PRIDE partner repository (https://www.ebi.ac.uk/pride/) (107) in MIAPE-compliant format with the dataset identifier PXD045458.

### Bacterial expression and purification of FBN proteins

Open reading frames of the mature sequences of AtFBN1a and AtFBN7a, without predicted N-terminal cTPs, were amplified by PCR and then cloned into pSMT3 vector providing an N-terminal 6X-His Tag followed by a SUMO tag. Details of restriction sites and primers are provided in **Table S3**. The plasmid constructs were transformed into BL21 (DE3) strain. An initial starter culture was grown at 37 ^0^C overnight and then 5 mL from the starter culture was added into 500 mL of the LB culture. The culture was induced using 0.1 mM IPTG (final concentration) at OD_600_ of 0.6 - 0.8 and incubated at 16 ^°^C or 22 ^°^C for 16 hours. The harvested cells were resuspended using the lysis buffer 50 mM HEPESLJKOH, pH 7.6, 250 mM NaCl, 10% glycerol, 0.5% Triton XLJ100, and 1 mM DTT and lysed using sonication. The lysed sample was centrifuged at 18000 rpm for 30 mins at 4 °C. Supernatant was filtered using a 0.45 µm nitrocellulose filter and loaded onto the Ni-NTA resin which was pre-equilibrated with Buffer A containing 50 mM HEPESLJKOH, pH 8.3, 250 mM NaCl, and 1 mM DTT. Consecutively, the resin was washed using Buffer A containing 20 mM, 40 mM, and 60 mM Imidazole. The tagged proteins were eluted using Buffer A containing 250 mM Imidazole. The eluants were dialyzed against the buffer containing 50 mM HEPESLJKOH, pH 8.3, 250 mM NaCl, 1 mM DTT, 10% (w/v) glycerol, 5% (w/v) glycerol and 1 mM EDTA. The proteins were concentrated using Amicon Ultra Centrifugal Filters (Millipore).

### P-96 fluorescence-based lipid binding assays

The lipid binding capability of FBNs was evaluated by monitoring the displacement of the fluorescently labelled fatty acid probe, P-96, i.e. 1-pyrenedodecanoic acid (108). The experiments were conducted with a Varian Cary Eclipse Fluorescence Spectrophotometer at a temperature of 25 °C in a quartz fluorometer cell (Starna Cells Inc., Atascadero, CA, USA). The excitation and emission wavelengths were set at 342 and 377 nm, respectively. We first estimated the K_d_ of P-96 with each protein variant by titrating increasing amounts of P-96 in a measurement buffer containing 10 mM MOPS at pH 7.2 (**Fig. S7**). Estimation of K_d_ was performed in the GraphPad Prism software v. 10.0.2.

Competition assays with each tested ligand were then performed using an amount ten times greater than the calculated K_d_ of P-96. The lipids used for the experiments were purchased from Sigma-Aldrich USA, Avanti Polar Lipids USA and Cayman Chemicals USA. To determine the binding capability, the fluorescence of P-96 was measured in two scenarios using a measurement buffer containing 175 mM mannitol, 0.5 mM K2SO4, 0.5 mM CaCl2, and 5 mM MES at pH 7.0, which was found to give a lower background fluorescence (109): firstly, without any proteins present (Q_0_), and secondly, with the addition of 0.5 µM of protein (Q_p_). Subsequently, P-96 was incubated with fatty acids for 1 minute before measuring fluorescence (F_0_). Then, 0.5 µM of protein was added, and the fluorescence was measured after an additional 2 minutes (F). The results were expressed as the percentage of fluorescence from the protein-P96 complex, calculated as (F-F_0_)/(Q_p_-Q_0_).

### Lipid-overlay assays

All the lipids were resuspended in chloroform or 100% ethanol to a final concentration of 5 mg/ml. Fatty acids were initially resuspended to 5 mg/mL and then serial dilution were performed. PVDF membrane strips were spotted with 0.75 µL of lipid and dried for 30 min at room temperature. Strips were blocked with TBS (Tris-HCl, pH 8.0, 150 mM NaCl) containing 3 % (w/v) lipid-free BSA for 1 h at room temperature. Then, strips were incubated with 100 nM (final concertation) of appropriate protein in TBST blocking solution (0.06% v/v Tween 20) at 4^0^ C for 16 h. Strips were washed 5 times for 10 min at room temperature with TBS with 0.1% (v/v) Tween 20 and then incubated with anti-His antibody conjugated with HRP (Rockland Immunochemicals, Inc, PA, USA) at 1:5000 dilution for 2 h at room temperature. Strips were washed 5 times for 10 min at room temperature with TBS with 0.1% (v/v) Tween-20 and then the bound protein was detected using Amersham ECL Western Blotting Detection Reagent (Cytiva, MA, USA) and visualized using Azure Biosystems (Dublin, CA, USA) C400 instrument.

### Circular dichroism

Protein concentrations were determined employing the Pierce^TM^ BCA protein quantification kit (Thermo Scientific, IL, USA). Subsequently, 300 µL of a 6 µM protein solution in a 10 mM phosphate buffer at pH 7.3 was introduced into a Hellama 1 mm QS cuvette, sealed with a PTFE stopper. A CD spectrum was then recorded using a JASCO-815 CD Spectrometer, with the wavelength range set between 190 and 260 nm. Data was collected in increments of 1 nm wavelength, with an averaging time of 1.5 seconds per wavelength point. Protein secondary structure was determined by analyzing the data derived from an average of five scans using the K2D3 server (http://cbdm-01.zdv.uni-mainz.de/∼andrade/k2d3/).

## DATA AVAILABILITY

All mass spectrometry proteomics data have been deposited at the ProteomeXchange Consortium via the PRIDE partner repository (https://www.ebi.ac.uk/pride/) in MIAPE-compliant format with the following unique identifier: PXD045458. Molecular Dynamics simulation data have been deposited with Zenodo at https://zenodo.org/record/8381676.

## SUPPORTING INFORMATION

This article contains supporting information.

### Supplemental Data File

HeliQuest: Amphipathic helix screening

### Supplemental Tables

Supplemental Table S1. Amphipathic Helix Predictions by HeliQuest

Supplemental Table S2. LC-MS/MS Proteomics of *N. benthamiana* Isolated Plastoglobules

Supplemental Table S3. List of primers used for cloning

### Supplemental Figures

Supplemental Figure S1. Plastoglobules hold a high proportion of proteins with predicted AHs

Supplemental Figure S2. Predicted amphipathic helices in *A. thaliana* Fibrillins

Supplemental Figure S3. Fibrillins show structural homology of Fibrillins and Seipins

Supplemental Figure S4. Confocal microscopy co-localization assays

Supplemental Figure S5. Initial simulation setups of each of the six replicas

Supplemental Figure S6. Lipid overlay assays of AtFBN1a and AtFBN7a

Supplemental Figure S7. P-96 binding curves

## ACKNOWLEDGEMENTS

We thank Doug Whitten of the MSU proteomics core facility for support in preparation and execution of proteomic analyses, Dr. Lisa Lapidus (MSU) for provision of the circular dichroism spectrometer, and Dr. Melinda Frame of the Center for Advanced Imaging at MSU for assistance in confocal microscopy analyses.

## AUTHOR CONTRIBUTIONS

P.K.L conceptualization; P.K.L data curation; K.-K.S., D.M.B. formal analysis; P.K.L funding acquisition; K.-K.S., D.M.B., A.H.-T. investigation; K.-K.S., J.V., P.K.L. methodology; P.K.L. project administration; D.M.B., J.V. software; J.V., P.K.L. supervision; P.K.L. visualization; K.-K.S., D.M.B., J.V., P.K.L. writing – original draft; K.-K.S., D.M.B., A.H.-T., J.V., P.K.L writing – review & editing.

## FUNDING AND ADDITIONAL INFORMATION

This work was supported by a grant from the National Science Foundation (MCB-2034631) to P.K.L. This work used Delta at NCSA through allocation BIO210061 from the Advanced Cyberinfrastructure Coordination Ecosystem Services & Supported (ACCESS) program, which is supported by National Science Foundation grants #2138259, #2138286. #2138307, #2137603, and #2138296 (110).

