## Supplemental Figures for "An amphipathic helix drives interaction of Fibrillins with plastoglobule lipid droplets"

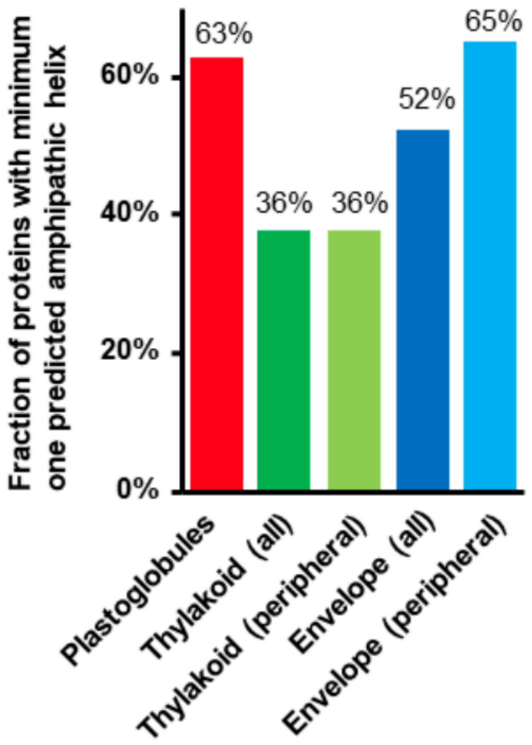


**Supplemental Figure S1. Plastoglobules hold a high proportion of proteins with predicted AHs.** Proteins with curated locations at select chloroplast sub-compartments were submitted to HeliQuest and analyzed for alpha-helical AHs using the same HeliQuest parameters.

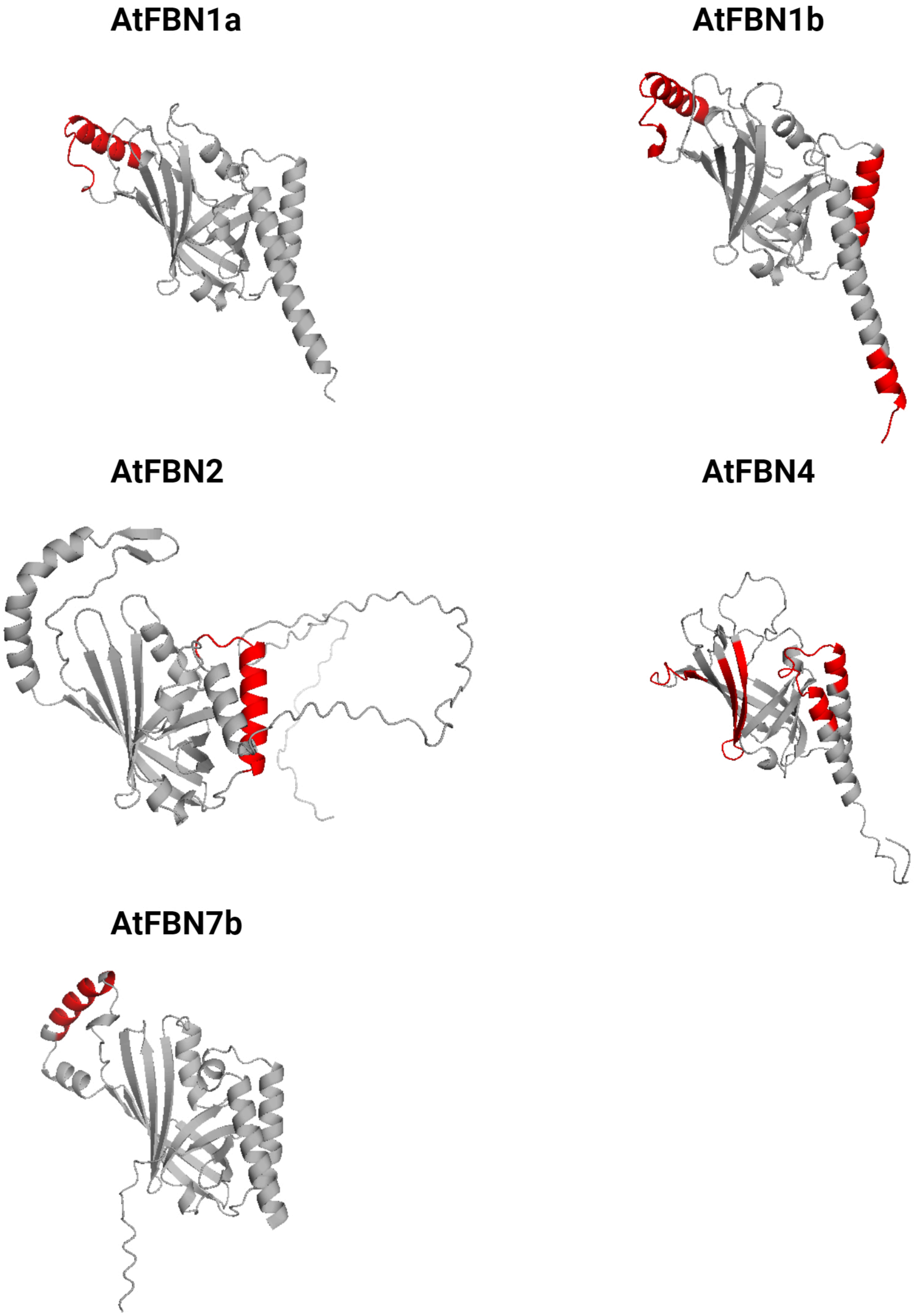


**Supplemental Figure S2. Predicted amphipathic helices in *A. thaliana* Fibrillins.** AlphaFold-predicted structures of each of the five FBN proteins with predicted amphipathic helices. Predicted amphipathic helices are highlighted in red.

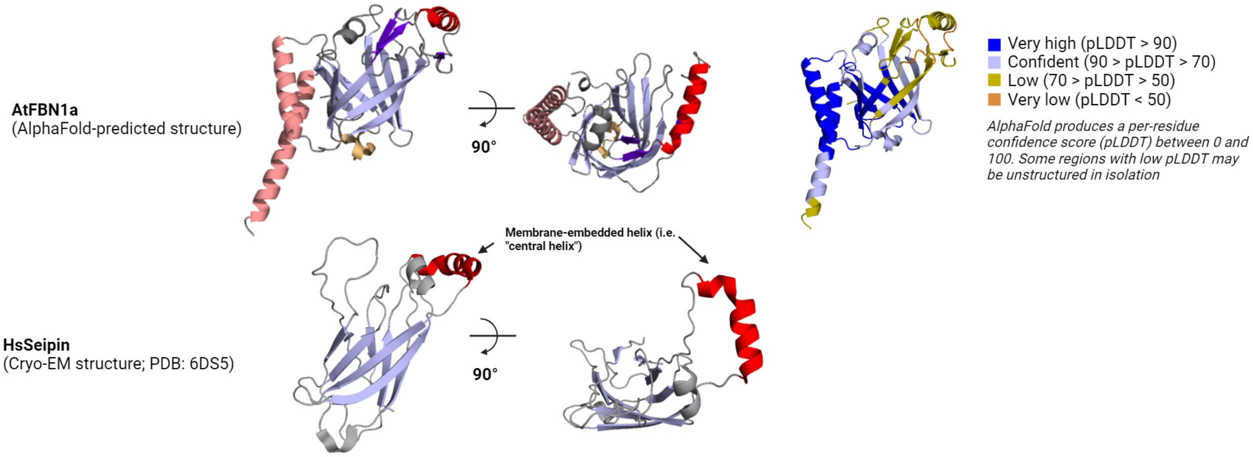


**Supplemental Figure S3. Fibrillins show structural homology of Fibrillins and Seipins.** Seipins are involved in Cyto-LD function and biogenesis.

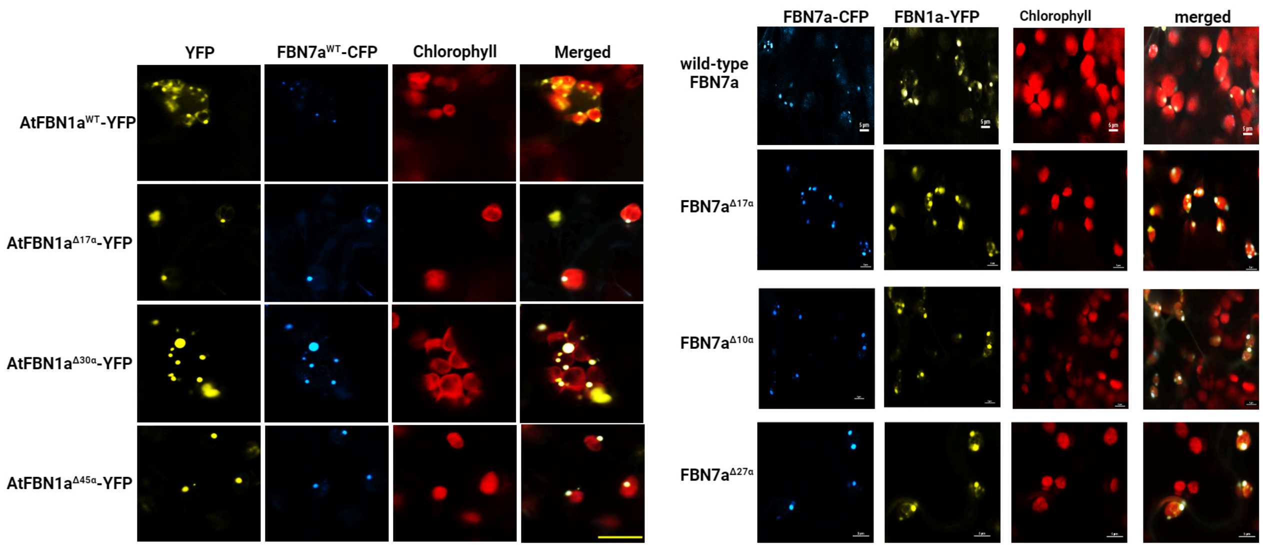


**Supplemental Figure S4. Confocal microscopy co-localization assays.** Tested constructs were tagged C-terminally with EYFP and a plastoglobule marker protein (AtFBN1a$^{WT}$ or AtFBN7a$^{WT}$) was tagged C-terminally with ECFP. Constructs were transiently co-transformed and observed by confocal microscopy 48 hours after transformation.

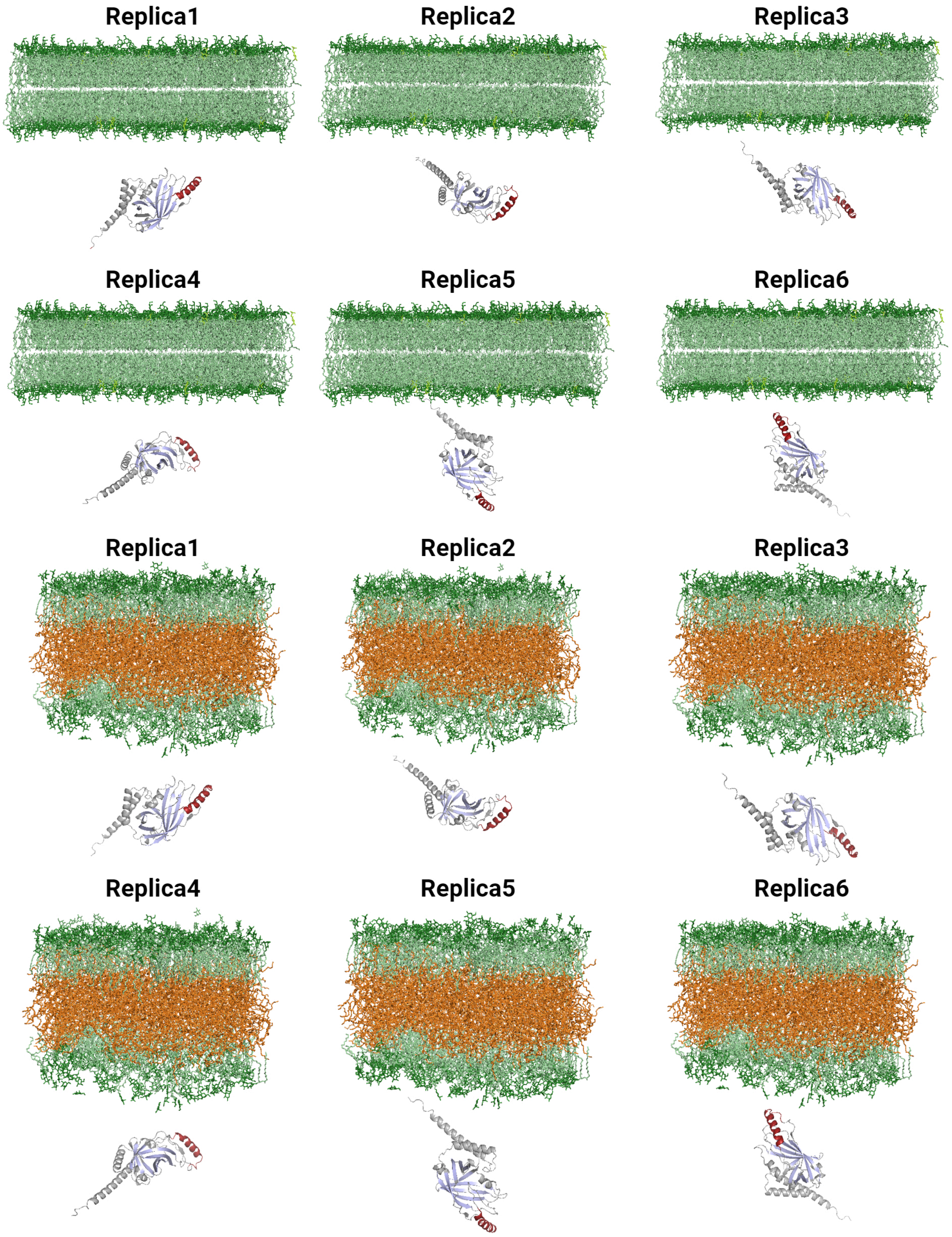


**Supplemental Figure S5. Initial simulation setups of each of the six replicas.** Plastoglobule trilayer (**A**), and the Thylakoid bilayer (**B**). Waters and ions are hidden. Coloring of membrane lipids and protein are as in the main text.

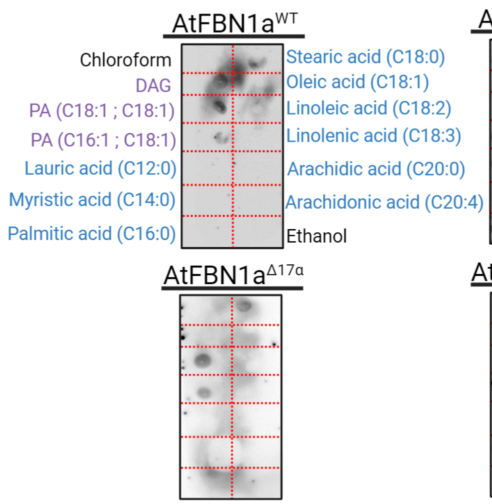


**Supplemental Figure S6. Lipid overlay assays of AtFBN1a and AtFBN7a.** Heterologously expressed and purified proteins (N-terminally tagged with His-SUMO) were incubated on PVDF membranes spotted with designated lipid and detected using anti-GFP antibody covalently linked to Horseradish Peroxidase. Position of lipids on the membranes is indicated for AtFBN1a$^{WT}$ and is unchanged for the other membranes.

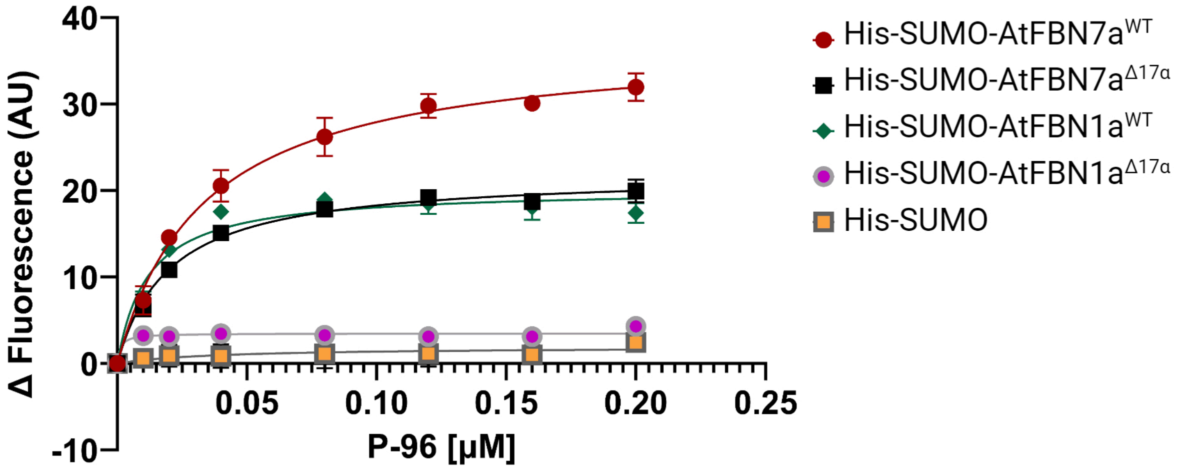


**Supplemental Figure S7. P-96 binding curves.** Each of the tested AtFBN1a and AtFBN7a variants and a His-SUMO construct as negative control are presented. Binding curves were used to estimate the dissociation constants, Kd using GraphPad Prism software.
