## Supplemental Data File for "An amphipathic helix drives interaction of Fibrillins with plastoglobule lipid droplets"

### HeliQuest: Amphipathic helix screening

**HeliQuest:**
**Alpha-helical Screen**

Database: PG

Size: 18

BlackList= No

Limit H: 0.3 <= H <= 0.6

Limit PRO= i, i+0 / N–0,N

Limit M: 0.35 <= M <= 1

Limit CYS= No

Limit Pol>= 0

Limit GLY>= 0

Limit Charg<= 8

Limit Res.Pol= nb mini= 0

Sum Charg: –3 <= Ch <= 3

Face H and Face P= Yes

Merged by protein and condensed plot

**ABC1K3 ...**

1 Seq LQAEARALGRAIDASIYS Pos 80 97

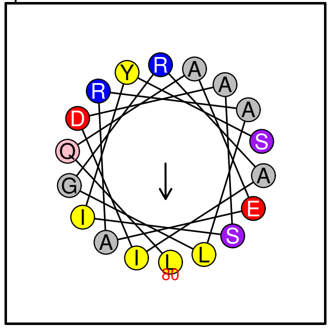

Helix

**ABC1K3 ...**

2 Seq LIRGVGKLINKYVDFITTDVLTLIDEF Pos 261 287

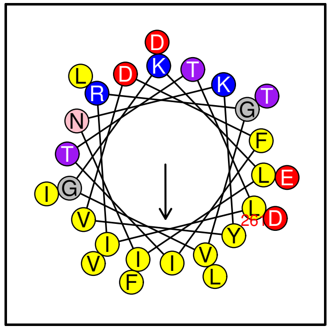

Helix/random coil

**ABC1K3 ...**

3 Seq AIIGHVVHLVNRDYEAMARDYYAL Pos 411 434

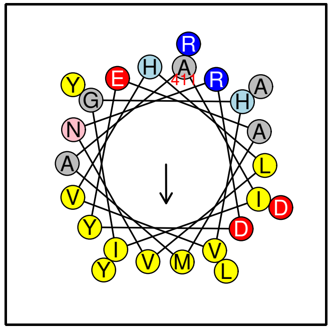

Helix/random coil

**ABC1K3 ...**

4 Seq AGGVGQRLAARFLQQLLRATT Pos 684 704

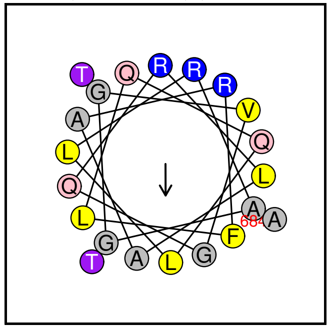

Possible lipid binding helix

**UnkSAG ...**

5 Seq QRIVSNLLENFISV../..LDGGSKILGMQDRL Pos 150 184

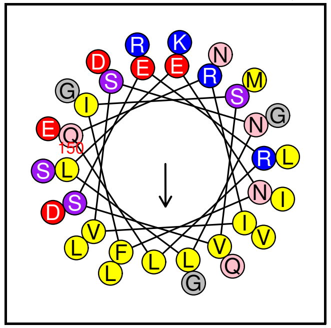

Helix

**UnkSAG ...**

6 Seq FKGMDSGLAGVTTLASTFD Pos 252 270

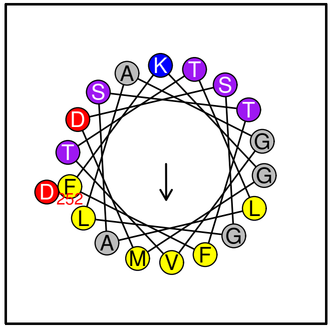

Helix/random coil

**ABC1K9 ...**

7 Seq RGVTRLVQGVQAFVGVGGEWLNDLSKS Pos 82 108

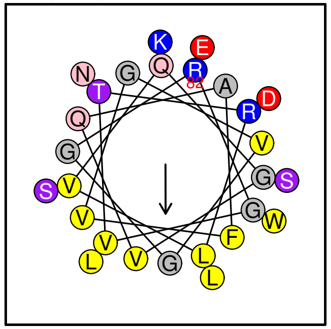

Possible lipid binding helix

**ABC1K9 ...**

8 Seq YLRKLFERMGATYIKLGQFIASA Pos 128 150

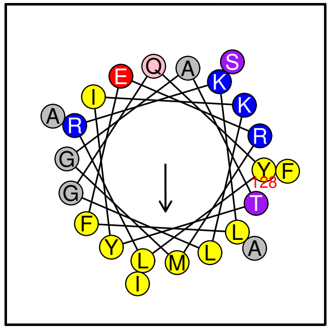

Lipid binding helix

**ABC1K9 ...**

9 Seq TSLVGIVKDIRESMLEEVDF Pos 258 277

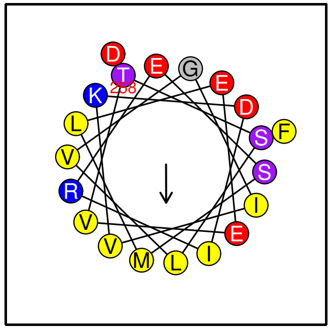

Helix/random coil

**ABC1K9 ...**
10 Seq AFAKDLEKMFSSIQELDTEIVVATAR Pos 428 453

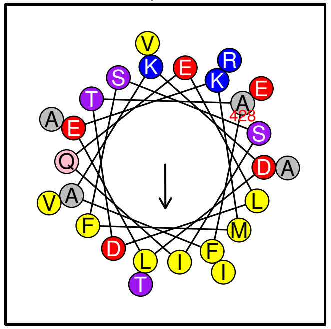

Helix

**ABC1K5 ...**
11 Seq ALDTLILRYIAGLI../..EWATSLFKEMDYLN Pos 229 271

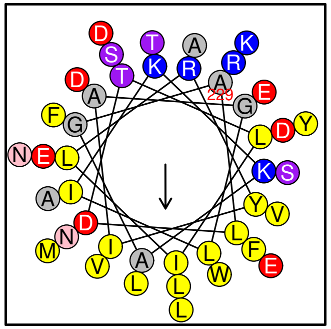

Helix

**ABC1K5 ...**
12 Seq LHLVNRDFKALAKDFVTLGLL Pos 381 401

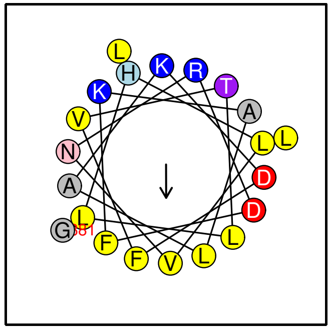

Helix/random coil

**ABC1K5 ...**
13 Seq AVTKALTDVFQDAISRGVRNISFG Pos 409 432

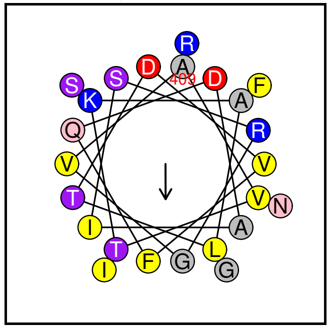

Helix

**ABC1K5 ...**
14 Seq LLREFAKGLDAYGLATLDSFT Pos 562 582

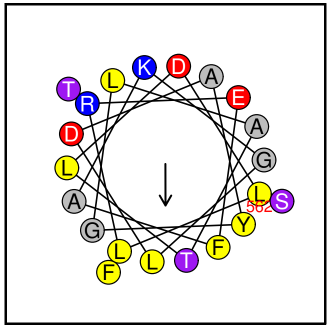

Helix/random coil

**LOX3 ...**
15 Seq TLVKHLDAFADKIGRNIV Pos 103 120

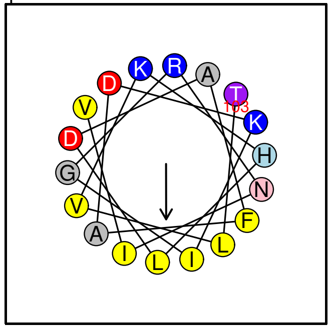

Possible lipid binding helix

**LOX3 ...**
16 Seq ELQSWYSESINVGHADLRD Pos 724 742

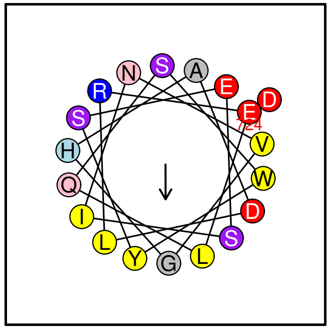

Helix/random coil

**FBN4 ...**
17 Seq KLLSVVSGLNRGLVASVDDLER Pos 90 111

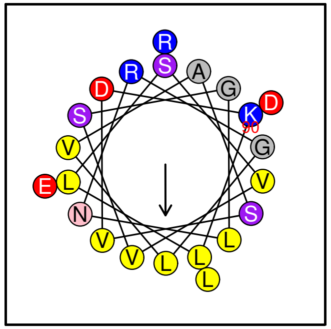

Helix/random coil

**FBN4 ...**
18 Seq TLGQVFQRIDVFSKDFDNIAEV Pos 168 189

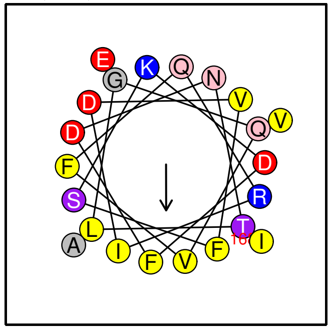

Helix/random coil

**ABC1K6 ...**
19 Seq KLSQSIESDVGELVNVGV Pos 386 403

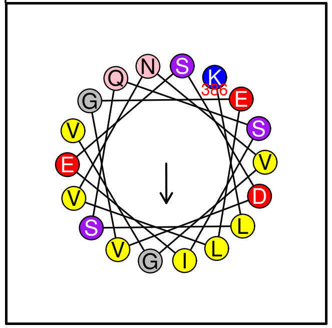

Helix/random coil

**ABC1K6 ...**
20 Seq YGMIEAIAHLIHRDYDAIVKDFVKLGFI Pos 450 477

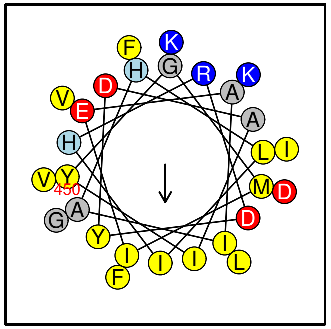

Helix/random coil

**ABC1K6 ...**
21 Seq DAERFIDVMQAFETFITAAKSGGG Pos 589 612

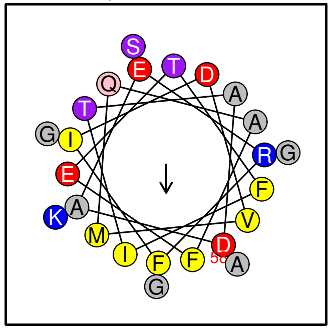

Helix

**ABC1K6 ...**
22 Seq FREFLLDEIVKGIDAITR Pos 664 681

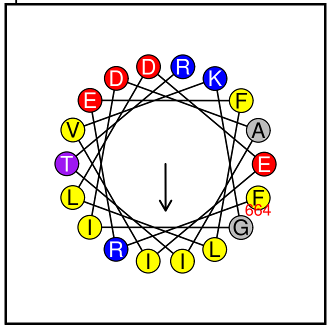

Helix/random coil

**ABC1K6 ...**
23 Seq EILSRLGSRVMARIVRDA Pos 774 791

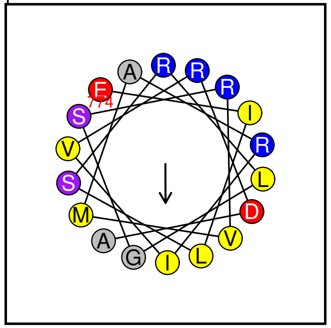

Possible lipid binding helix

**PLIN4 ...**
24 Seq DAVSSGVASVVDVAKGVVQGGLDT Pos 90 113

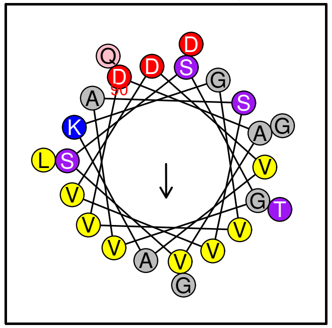

Helix/random coil

**PLIN4 ...**
25 Seq GLLRQLHTAYSGLVSSLQGL Pos 1261 1280

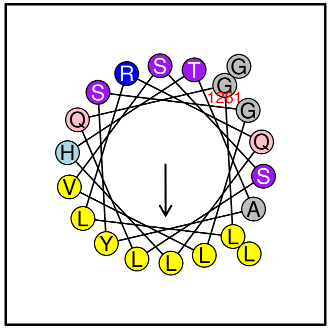

Possible lipid binding helix

**PLIN4 ...**
26 Seq REGVHQAWQGLEQLLEGLQHN Pos 1320 1340

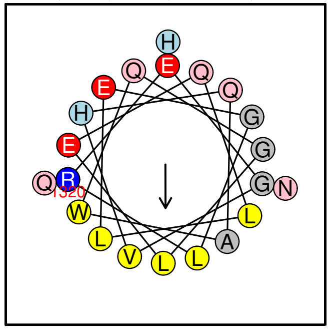

Helix

**Unk1 ...**
27 Seq LGKALAEVINERIESTVGEVLSTIGKFQAE Pos 78 107

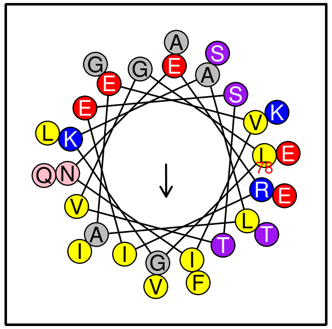

Helix

**ArfGAP1 ...**

28 Seq DDFLNNAMSSLYSGWSSFTTGASRFASA Pos 197 224

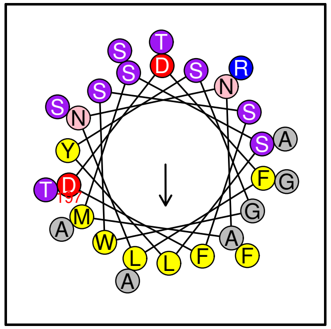

Helix/random coil

**ArfGAP1 ...**

29 Seq IFDDVSSGVSQLAS../..GSKGWRDVTTFFSG Pos 264 296

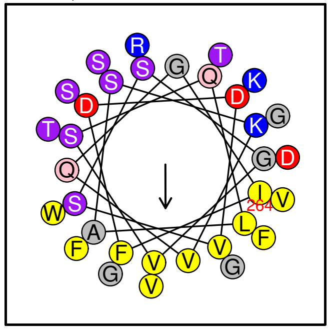

Helix/random coil

**LOX4 ...**

30 Seq TLVKHLDAFTDKIGRNVVL Pos 109 127

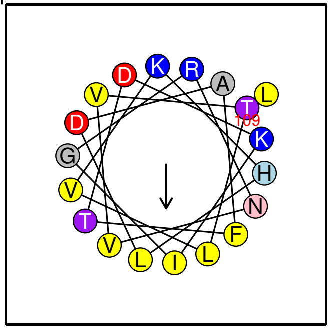

Possible lipid binding helix

**LOX4 ...**

31 Seq WSAIQTWVRTYVERYYAN Pos 705 722

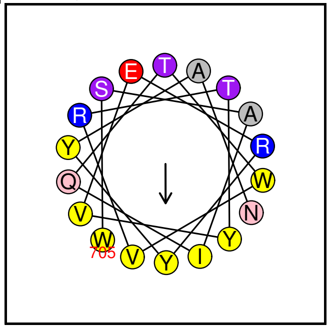

Possible lipid binding helix

**FBA–iso2 ...**

32 Seq HHGIDRTYDVAEKVWAEVF Pos 234 252

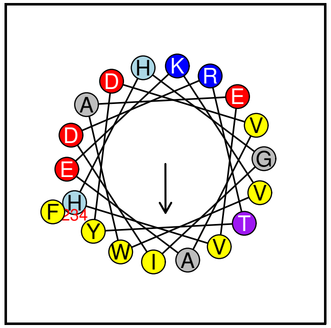

Helix/random coil

**FBA2–iso1 ...**

33 Seq HDIDRTYDVAEKVWAEVF Pos 234 251

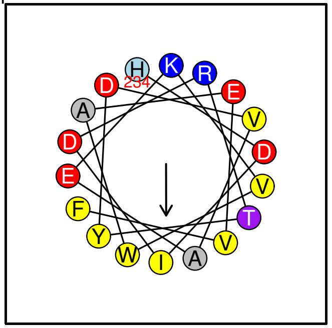

Helix/random coil

**FBA–iso3 ...**

34 Seq HGIDRTYDVAEKVWAEVF Pos 235 252

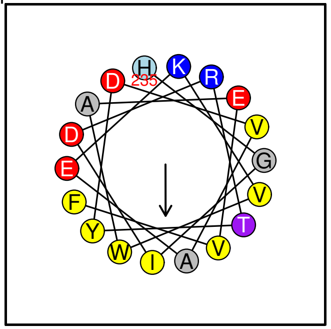

Helix/random coil

**FBA1–iso1 ...**

35 Seq HGIDRTYDVAEKVWAEVF Pos 235 252

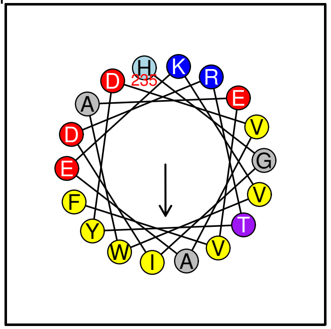

Helix/random coil

**GMAP–210 ...**

36 Seq MSSWLGGLGSGLGQ../..SLTGQISNFTKDML Pos 1 38

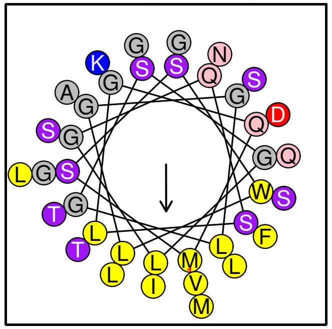

Helix/random coil

**GMAP–210 ...**
37 Seq LSNEVSRLESEVGHWRHI Pos 177 194

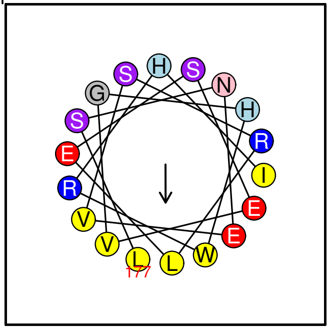

Helix/random coil

**GMAP–210 ...**
38 Seq LQQALSDAENEIMRLSSL Pos 390 407

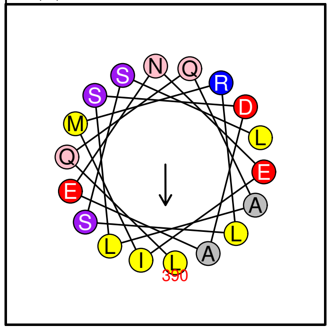

Helix

**GMAP–210 ...**
39 Seq LKNFGQELAQVQHSIGQL Pos 1281 1298

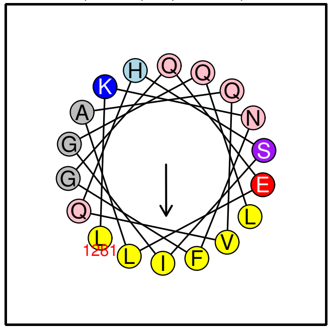

Helix

**GMAP–210 ...**
40 Seq NELLRQAVTNLKERILILE Pos 1437 1455

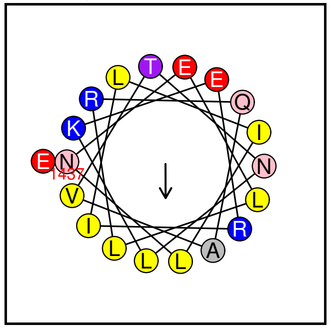

Helix/random coil

**GMAP–210 ...**
41 Seq GGVTRWMTGWLGGGSKSV Pos 1826 1843

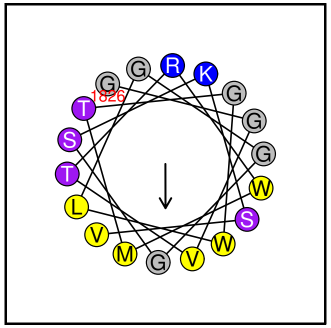

Possible lipid binding helix

**ABC1K1–iso2 ...**
42 Seq WKRLENLLSLAKENVAKM Pos 544 561

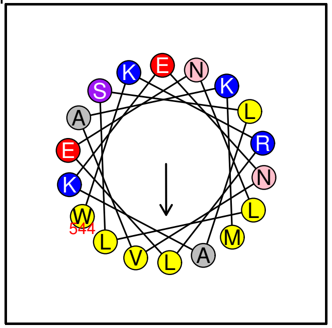

Possible lipid binding helix

**ABC1K1–iso1 ...**
43 Seq WKRLENLLSLAKENVAKM Pos 569 586

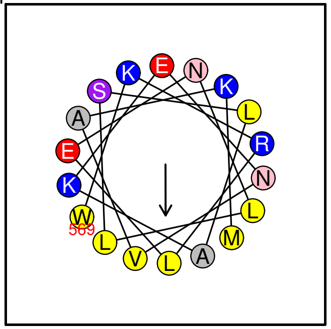

Possible lipid binding helix

**PEX11 ...**
44 Seq RDKLLRTIQYFSRFYAWY Pos 25 42

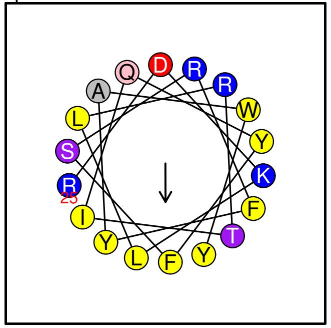

Possible lipid binding helix

**PEX11 ...**
45 Seq RIGKFLEHLKAAAVAFDNK Pos 71 89

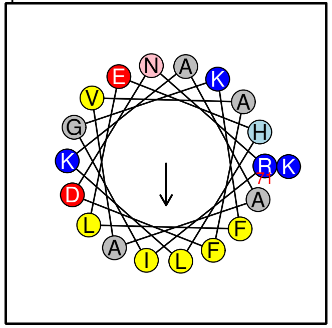

Possible lipid binding helix

**PEX11 ...**
46 Seq GILNLDDGIVGITG../..SLIGVWSQWRKTAG Pos 206 236

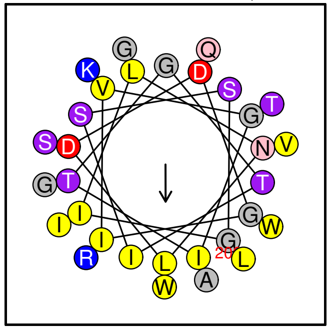

Helix

**AOS ...**
47 Seq FQATYSELFDSLEKELSL Pos 186 203

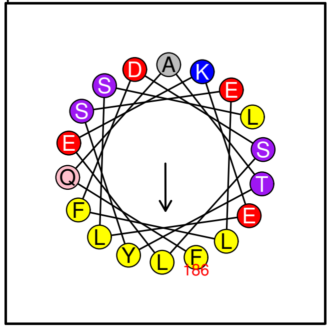

Helix

**AOS ...**
48 Seq LVKSDYQRLYEFFLESAGEIL Pos 277 297

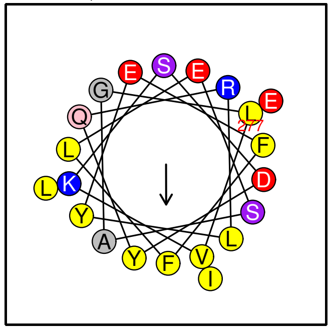

Helix

**LOX2–iso2 ...**
49 Seq RIIKALGEAQDDILQFDA Pos 224 241

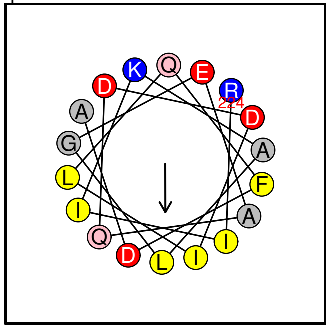

Helix/random coil

**LOX2–iso2 ...**
50 Seq VINAAFERFKGKLQYLEGVID Pos 693 713

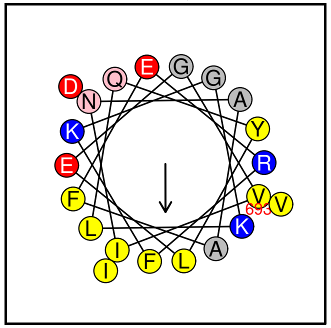

Helix/random coil

**LOX2–iso1 ...**
51 Seq RIIKALGEAQDDILQFDA Pos 366 383

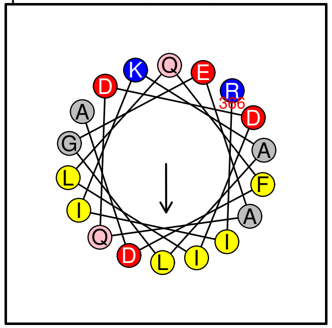

Helix/random coil

**LOX2–iso1 ...**
52 Seq VINAAFERFKGKLQYLEGVID Pos 835 855

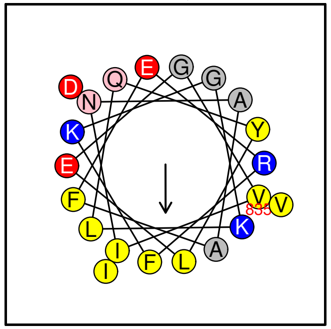

Helix/random coil

**FBN1a ...**
53 Seq IKGLLTSVQDTASSVARTISN Pos 249 269

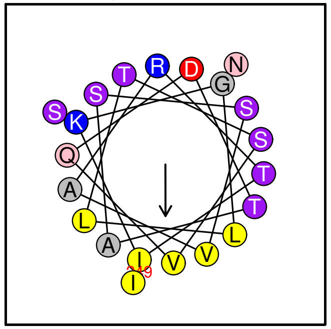

Possible lipid binding helix

**FBN1b ...**
54 Seq IRGLLTSVQDTASSVARTISS Pos 241 261

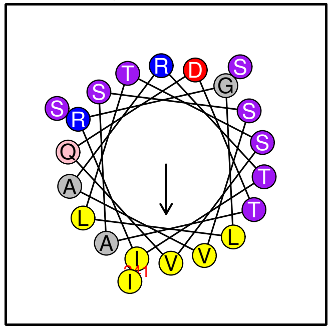

Possible lipid binding helix

**PLIN1 ...**
55 Seq SGGADLALGSIEKVVEYL Pos 174 191

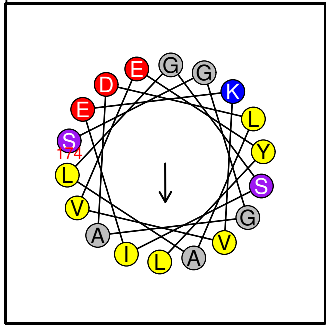

Helix/random coil

**PLIN1 ...**
56 Seq GLLGGVAHTLQKTLQTTISAVT Pos 330 351

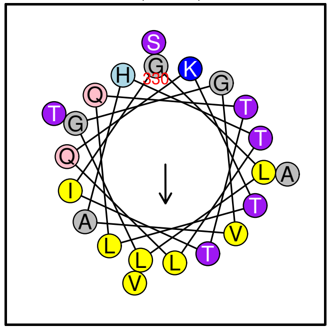

Possible lipid binding helix

**PLIN1 ...**
57 Seq AMSLSDALKGVTDNVVDTVVHYV Pos 380 402

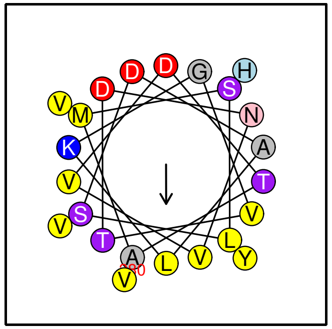

Helix/random coil

**M48−iso2 ...**
58 Seq GLNEFGKALLGSMTEQIMLLENIG Pos 94 117

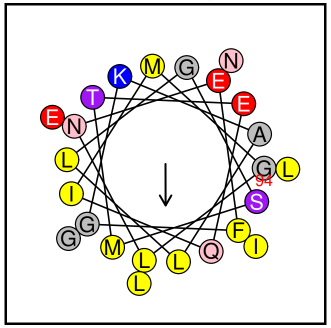

Helix/random coil

**M48−iso2 ...**
59 Seq KNQLSDLHGLLVEAAEILN Pos 124 142

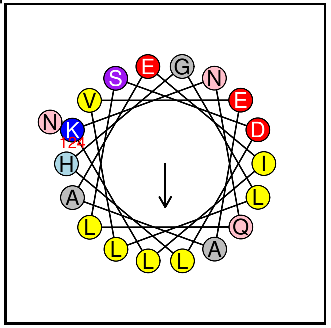

Helix

**M48−iso1 ...**
60 Seq GLNEFGKALLGSMTEQIMLLENIG Pos 94 117

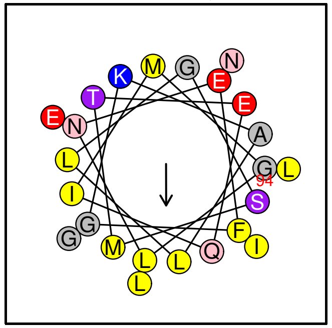

Helix/random coil

**M48−iso1 ...**
61 Seq KNQLSDLHGLLVEAAEILN Pos 124 142

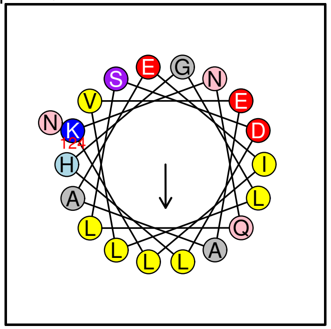

Helix

**NDC1 ...**
62 Seq FAKSAVDSIALLQSNLTKVLSG Pos 497 518

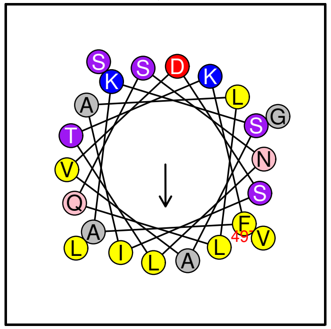

Possible lipid binding helix

**ABC1K7−iso3 ...**
63 Seq SWADENYSSLQRSIDVWS Pos 138 155

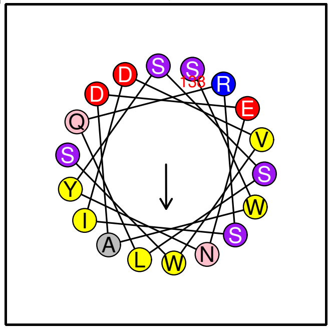

Helix/random coil

**ABC1K7-iso2 ...**
64 Seq SWADENYSSLQRSIDVWS Pos 138 155

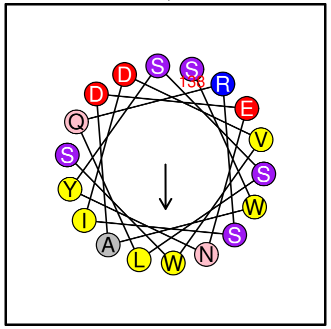

Helix/random coil

**ABC1K7-iso1 ...**
65 Seq SWADENYSSLQRSIDVWS Pos 138 155

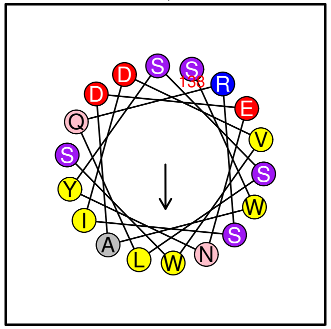

Helix/random coil

**PLIN3 ...**
66 Seq QLSQVLSLMETVKQGVDQ Pos 271 288

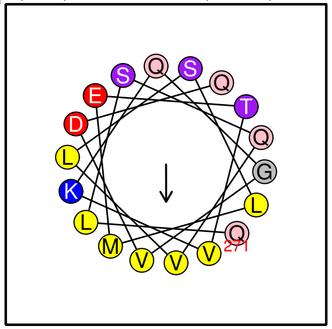

Helix

**PLIN3 ...**
67 Seq EQVESRALTMFRDIAQQLQA Pos 320 339

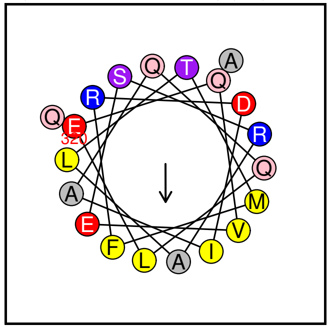

Helix

**PPH-iso3 ...**
68 Seq YTYWRGVRESFRSSFIRVFGG Pos 484 504

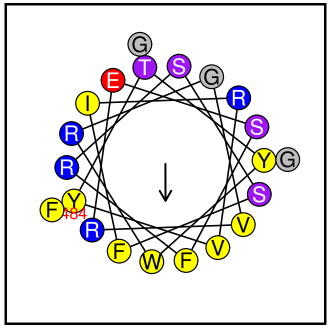

Possible lipid binding helix

**PPH-iso2 ...**
69 Seq YTYWRGVRESFRSSFIRVFGG Pos 461 481

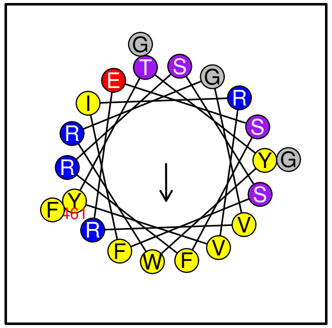

Possible lipid binding helix

**PPH-iso1 ...**
70 Seq YTYWRGVRESFRSSFIRVFGG Pos 461 481

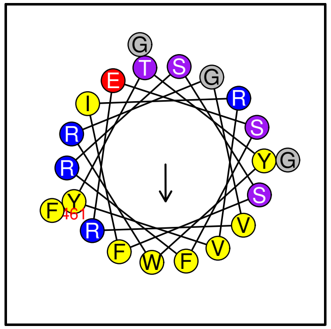

Possible lipid binding helix

**FBN2 ...**
71 Seq FKAGSEVRAEVLELVNQLEAL Pos 168 188

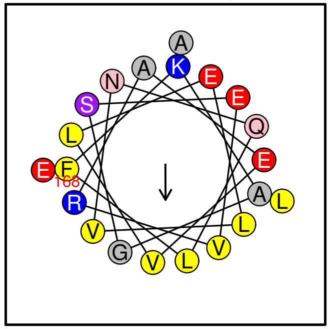

Helix

**PES1 ...**
72 Seq KGELAYALDEVLGFLRNA Pos 403 420

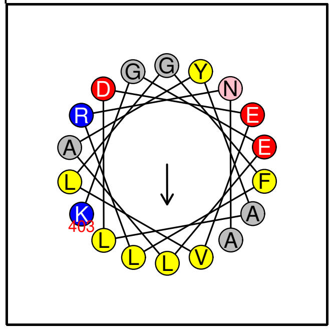

Helix

**PES1 ...**
73 Seq VYLEVKAEVENSIAYLLKK Pos 656 674

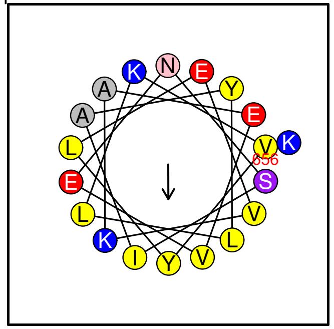

Helix/random coil

**PES2 ...**
74 Seq SVQRMGGVGGGMLRDVLAV Pos 269 287

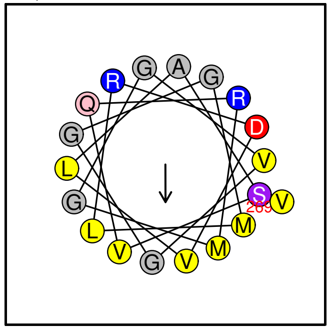

Helix/random coil

**AKR ...**
75 Seq YLDGLGDAVEQGLVKAVG Pos 189 206

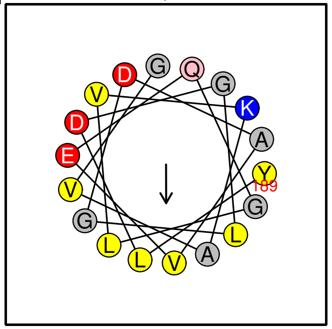

Helix/random coil

**Est1 ...**
76 Seq FLEDGVDLVSIIKRAYYY Pos 347 364

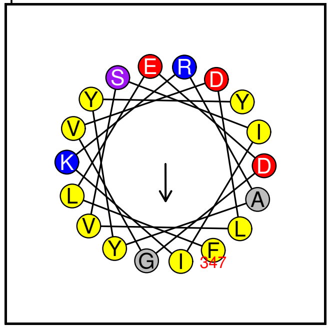

Helix/random coil

**CCD4 ...**
77 Seq NVFSGFNGVTASVARGAL Pos 201 218

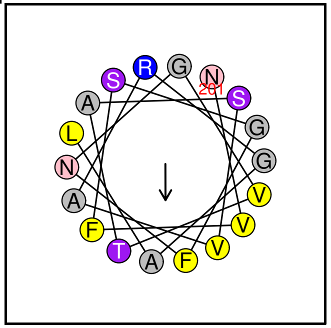

Helix/random coil

**FRR2 ...**
78 Seq VAGATGQTGKRIVEQLLS Pos 51 68

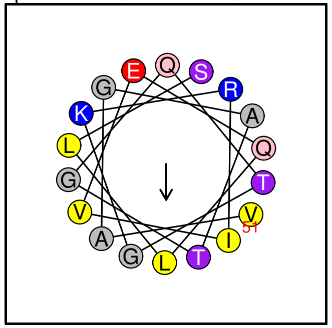

Helix/random coil

**Fred2 ...**
79 Seq VAGATGQTGKRIVEQLLS Pos 51 68

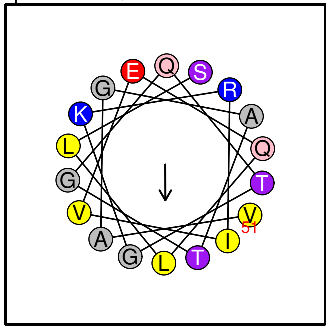

Helix/random coil

### HeliQuest: Amphipathic helix screening

**HeliQuest:**
**3-11 Helical Screen**

Database: PG

Size: 11

BlackList= No

Limit H: 0.27 <= H <= 0.35

Limit PRO= i, i+0 / N–0,N

Limit M: 0.27 <= M <= 0.45

Limit CYS= No

Limit Pol>= 5

Limit GLY>= 1

Limit Charg: 0 <= nCh <= 4

Net Charg: –3 <= z <= 3

Limit Res.Hyd= TAILV nb mini= 5

Face H = Yes

Merged by protein and condensed plot

Face H and Face P= No

**LOX2−iso2 ...**

1 Seq EFARQTLAGLN Pos 257 267

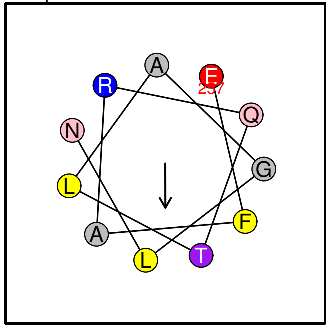


No prediction (3−11 type)

**LOX2−iso1 ...**

2 Seq EFARQTLAGLN Pos 399 409

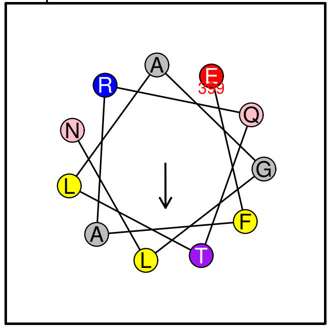


No prediction (3−11 type)

**FBN1b ...**

3 Seq ATDTGEIGSAL Pos 60 70

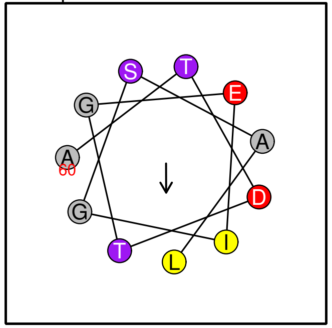


No prediction (3−11 type)

**FBN1b ...**

4 Seq ETRAEIGDLITQ Pos 106 117

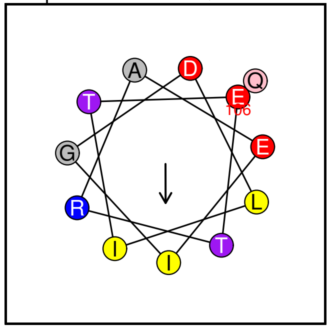


No prediction (3−11 type)

**FBN1b ...**

5 Seq RGLLTSVQDTA Pos 242 252

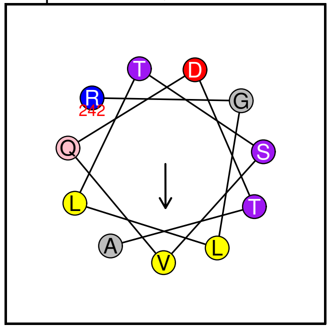


No prediction (3−11 type)

**FBN1a ...**

6 Seq KGLLTSVQDTA Pos 250 260

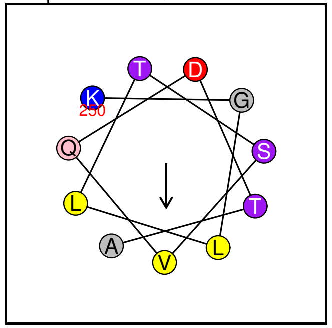


No prediction (3−11 type)

**AKR ...**

7 Seq DAVEQGLVKAV Pos 195 205

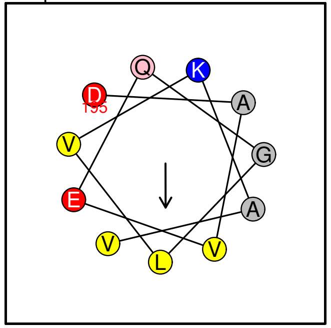


No prediction (3−11 type)

**PLIN1 ...**

8 Seq IASTSDKVLGAALAG Pos 135 149

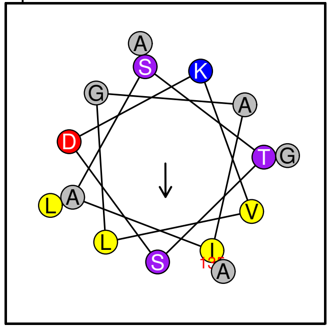


No prediction (3−11 type)

**PLIN1 ...**

9 Seq GADLALGSIEK Pos 176 186

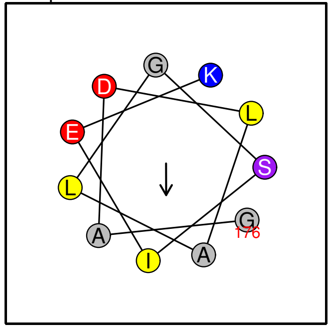


No prediction (3−11 type)

**PLIN1 ...**

10 Seq SRVGALTNTLS Pos 220 230

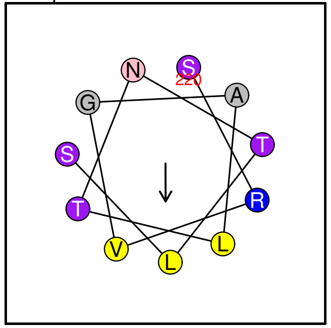

No prediction (3–11 type)

**PLIN1 ...**

11 Seq MARALEQGHTV Pos 237 247

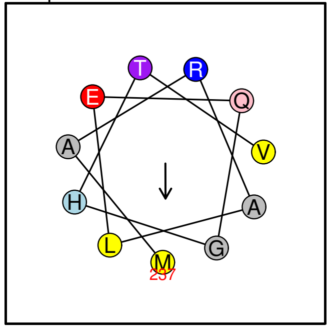

No prediction (3–11 type)

**ABC1K7–iso3 ...**

12 Seq VLHNGEKVVVKV Pos 277 288

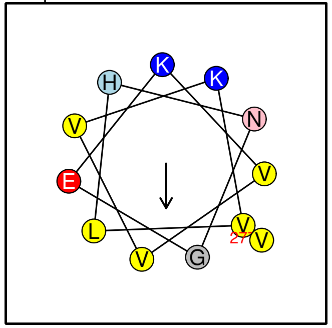

No prediction (3–11 type)

**ABC1K7–iso3 ...**

13 Seq INNLDALAARGFN Pos 384 396

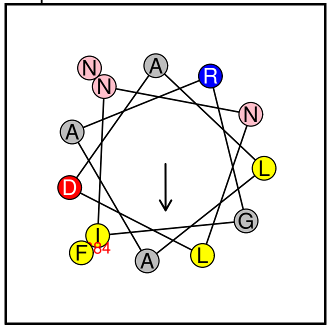

No prediction (3–11 type)

**ABC1K7–iso2 ...**

14 Seq VLHNGEKVVVKV Pos 277 288

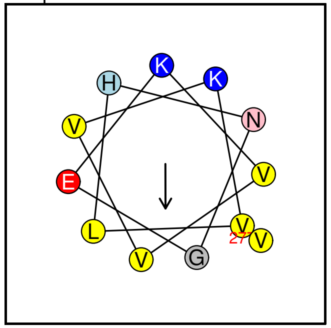

No prediction (3–11 type)

**ABC1K7–iso2 ...**

15 Seq INNLDALAARGFN Pos 384 396

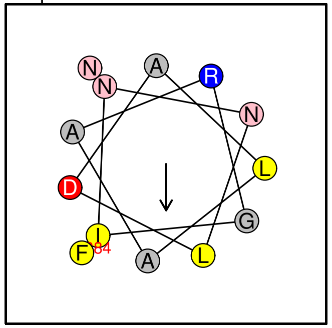

No prediction (3–11 type)

**ABC1K7–iso1 ...**

16 Seq VLHNGEKVVVKV Pos 277 288

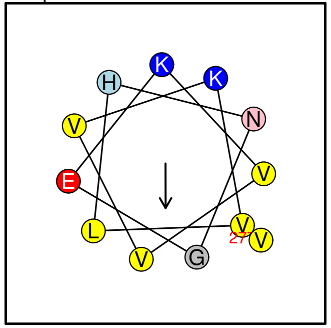

No prediction (3–11 type)

**ABC1K7–iso1 ...**

17 Seq INNLDALAARGFN Pos 384 396

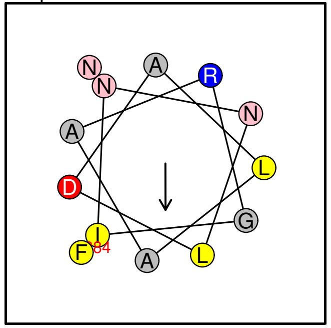

No prediction (3–11 type)

**ClpR1 ...**

18 Seq IELLAKGTGKS Pos 313 323

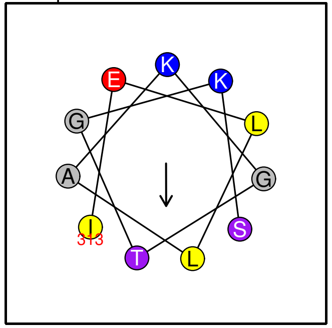

No prediction (3–11 type)

**PLIN4 ...**
19 Seq SGVASVVDVAKGVVQGGLD Pos 94 112

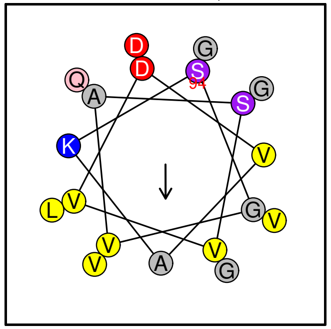

No prediction (3–11 type)

**PLIN4 ...**
20 Seq SALTGTKEVVSS Pos 116 127

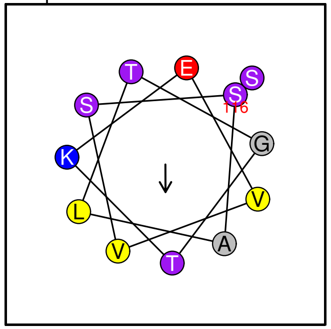

No prediction (3–11 type)

**PLIN4 ...**
21 Seq QGGLDTSKAVLTGTKDTVST Pos 141 160

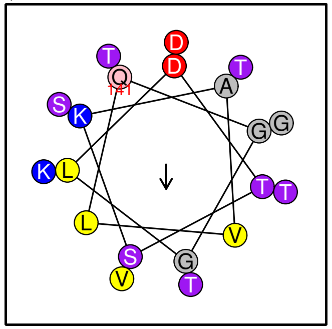

No prediction (3–11 type)

**PLIN4 ...**
22 Seq LTGAVNVAKGTVQA../..TKTVLTGTKDTVTT Pos 162 193

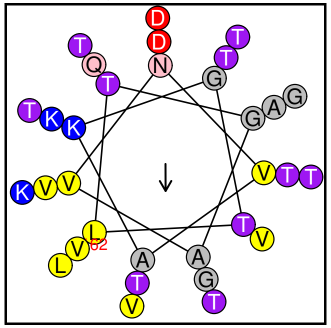

No prediction (3–11 type)

**PLIN4 ...**
23 Seq GAVNLAKGTVQT Pos 197 208

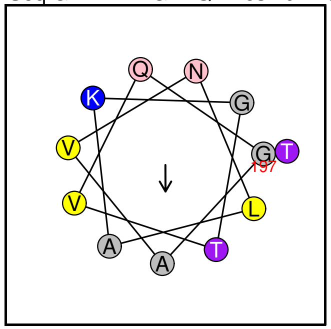

No prediction (3–11 type)

**PLIN4 ...**
24 Seq AVLTGTKDAVST Pos 215 226

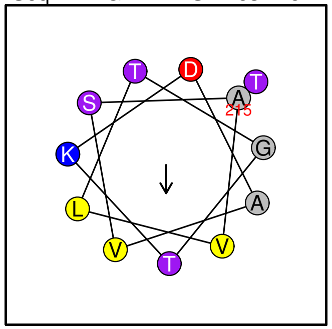

No prediction (3–11 type)

**PLIN4 ...**
25 Seq GAVNVARGSIQTGVDTSKTVLTGT Pos 230 253

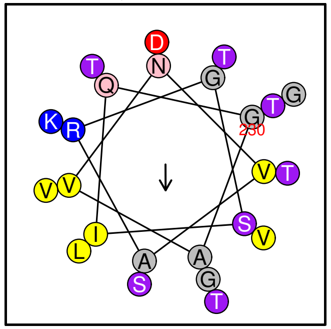

No prediction (3–11 type)

**PLIN4 ...**
26 Seq TGAMNVAKGTIQTGVDTSKTVLTGT Pos 262 286

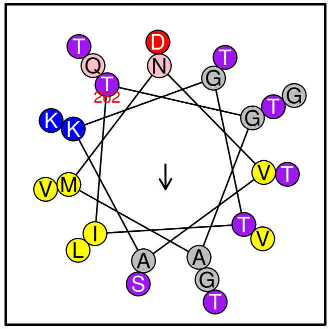

No prediction (3–11 type)

**PLIN4 ...**
27 Seq TGAMNVAKGTIQTGVDTSKTVLTGT Pos 295 319

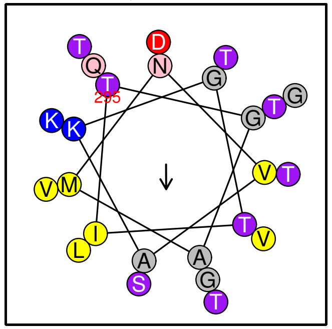

No prediction (3–11 type)

**PLIN4 ...**
28 Seq TGAMNVAKGTIQTGVDTTKTVLTGT Pos 328 352

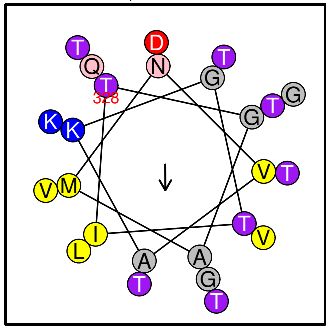

No prediction (3–11 type)

**PLIN4 ...**
29 Seq VTGAVNLAKEAIQGGLD Pos 360 376

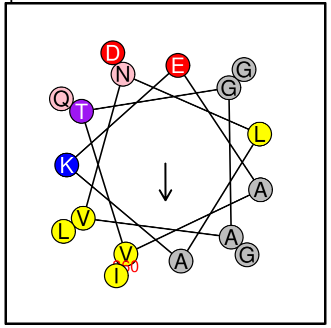

No prediction (3–11 type)

**PLIN4 ...**
30 Seq DTMSTGLTGAAN Pos 387 398

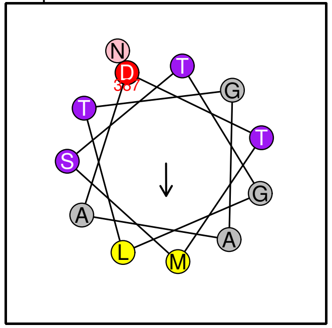

No prediction (3–11 type)

**PLIN4 ...**
31 Seq LNTTQNIATGT Pos 408 418

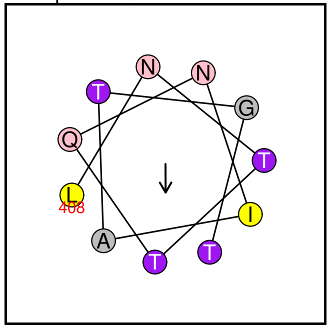

No prediction (3–11 type)

**PLIN4 ...**
32 Seq GAMNLARGTIQTGVDTTKIVLTGTKDT Pos 428 454

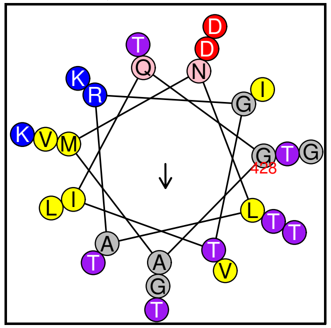

No prediction (3–11 type)

**PLIN4 ...**
33 Seq QGGLDTTKSVLTGTKDAVST Pos 471 490

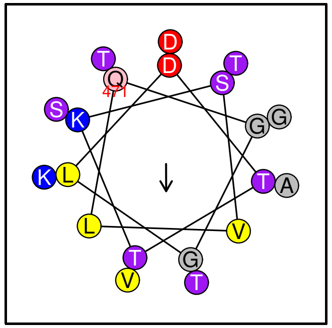

No prediction (3–11 type)

**PLIN4 ...**
34 Seq LTGAVNVAKGTVQTGVDTTKTVLTGT Pos 492 517

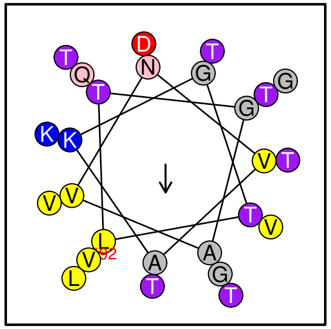

No prediction (3–11 type)

**PLIN4 ...**
35 Seq DTMSTGLTGAAN Pos 552 563

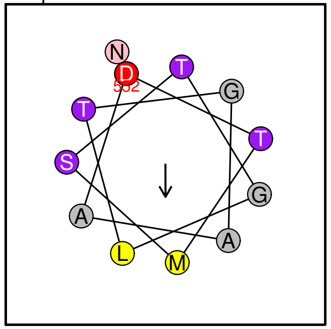

No prediction (3–11 type)

**PLIN4 ...**
36 Seq QTGVDTAKTVLTGTKDTVTT Pos 570 589

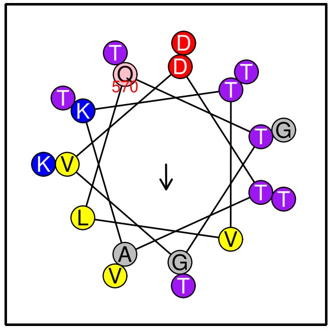

No prediction (3–11 type)

**PLIN4 ...**
37 Seq AVNVAKGTVQTGMD../..TQNIATGTKNTFGS Pos 594 655

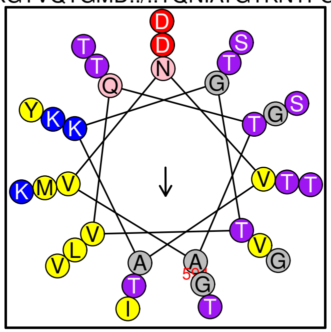

No prediction (3−11 type)

**PLIN4 ...**
38 Seq QTGVDTAKTVLTGTKDTVTT Pos 669 688

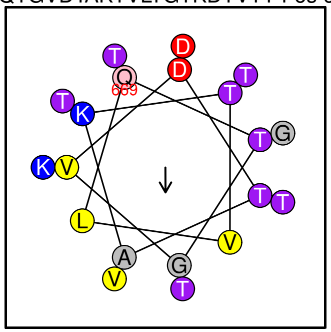

No prediction (3−11 type)

**PLIN4 ...**
39 Seq AVNVAKGTVQT Pos 693 703

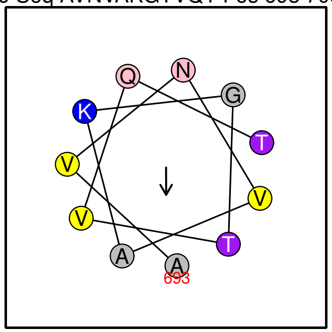

No prediction (3−11 type)

**PLIN4 ...**
40 Seq SVDTTKTVLTGT Pos 704 715

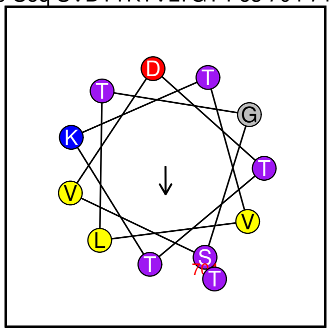

No prediction (3−11 type)

**PLIN4 ...**
41 Seq NVAKGAIQGGLDTT../..QTGMDTTKTVLTGT Pos 728 781

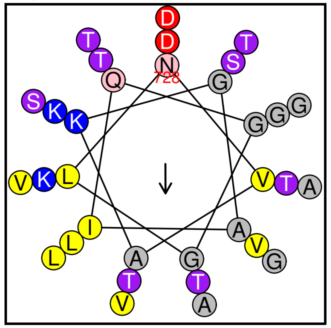

No prediction (3−11 type)

**PLIN4 ...**
42 Seq VQMGVDTAKTVLTGT Pos 800 814

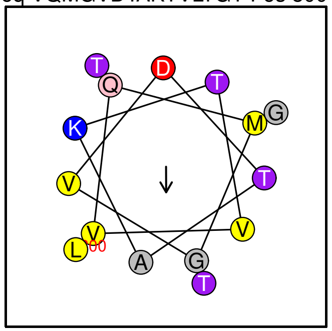

No prediction (3−11 type)

**PLIN4 ...**
43 Seq VQTGLKTTQNIATGT Pos 833 847

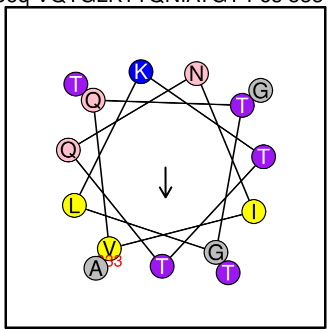

No prediction (3−11 type)

**PLIN4 ...**
44 Seq NTLGSGVTGAAK Pos 849 860

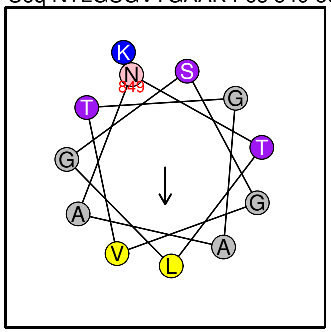

No prediction (3−11 type)

**PLIN4 ...**
45 Seq QGGLDTTKSVLTGTKDAVST Pos 867 886

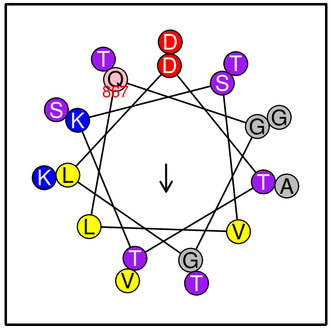

No prediction (3−11 type)

**PLIN4 ...**

46 Seq GAVNLAKGTVQTGVDTSKTVLTGT Pos 890 913

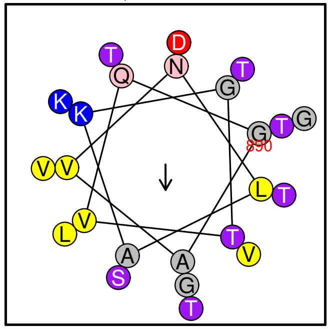

No prediction (3–11 type)

**PLIN4 ...**

47 Seq AVNVAKGTVQTGVDTAKTVLSGAKDAVTT Pos 924 952

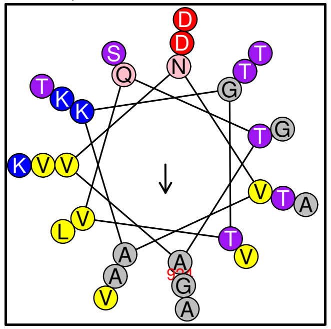

No prediction (3–11 type)

**PLIN4 ...**

48 Seq AVNVAKGTVQTGVDASKAVLMGTKDTV Pos 957 983

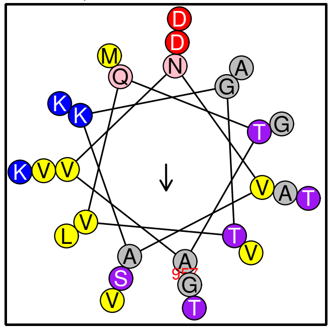

No prediction (3–11 type)

**PLIN4 ...**

49 Seq QGGLDTTKTVLTGTKDAVSA Pos 999 1018

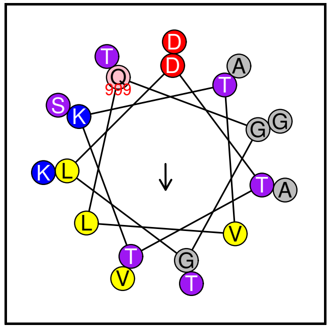

No prediction (3–11 type)

**VTE1 ...**

50 Seq NVFEGATGEVA Pos 308 318

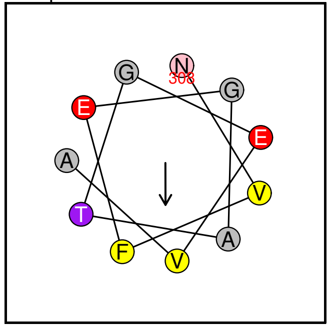

No prediction (3–11 type)

**Unk1 ...**

51 Seq VINERIESTVG Pos 85 95

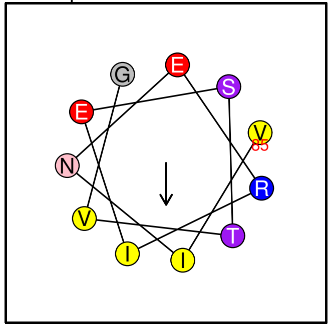

No prediction (3–11 type)

**LOX4 ...**

52 Seq DKIGRNVVLEL Pos 119 129

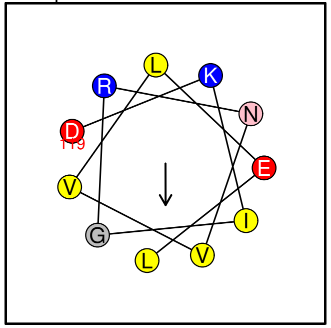

No prediction (3–11 type)

**GMAP–210 ...**

53 Seq DNSAGVVLKDL Pos 1966 1976

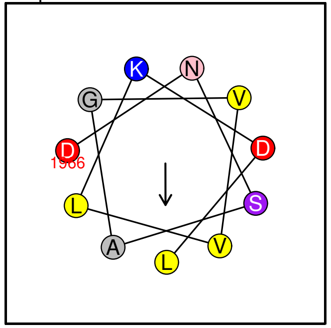

No prediction (3–11 type)

**FBN7b–iso4 ...**

54 Seq LRAMVQETVQG Pos 47 57

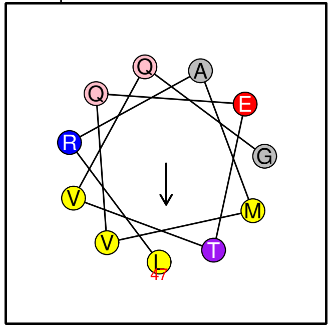

No prediction (3–11 type)

**FBN7b–iso4 ...**
55 Seq QLKSALGQAATTL Pos 220 232

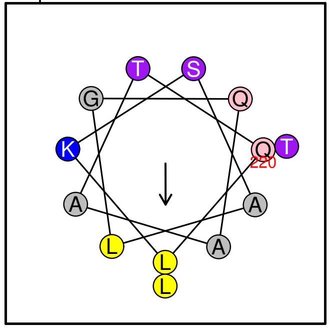

No prediction (3–11 type)

**FBN7b–iso2 ...**
56 Seq LRAMVQETVQG Pos 47 57

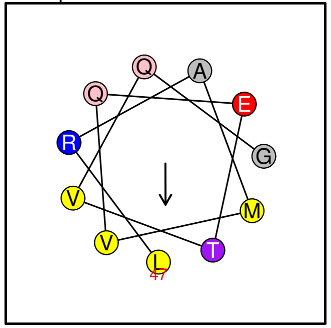

No prediction (3–11 type)

**FBN7b–iso2 ...**
57 Seq QLKSALGQAATTL Pos 220 232

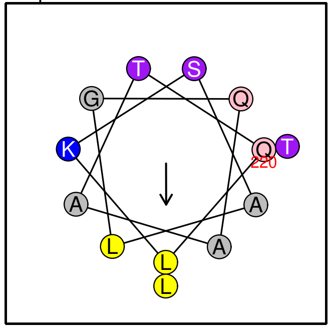

No prediction (3–11 type)

**FBN7b–iso1 ...**
58 Seq LRAMVQETVQG Pos 47 57

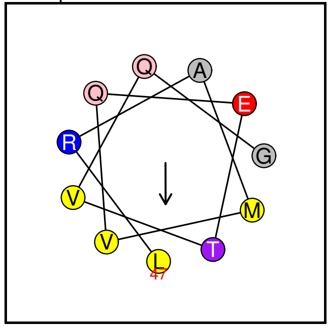

No prediction (3–11 type)

**FBN7b–iso1 ...**
59 Seq QLKSALGQAATTL Pos 220 232

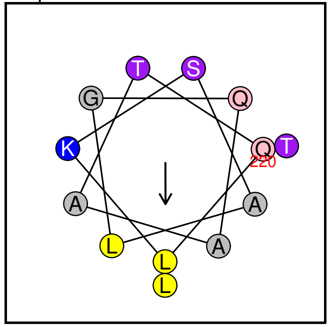

No prediction (3–11 type)

**ArfGAP1 ...**
60 Seq FRDKVVALAEG Pos 111 121

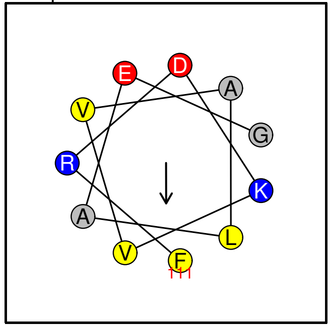

No prediction (3–11 type)

**FBA2–iso2 ...**
61 Seq TEGKKMVDVLV Pos 127 137

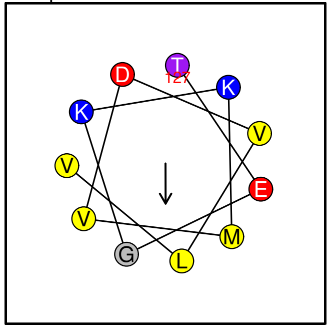

No prediction (3–11 type)

**FBA2–iso1 ...**
62 Seq TEGKKMVDVLV Pos 127 137

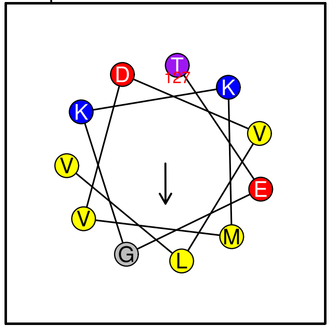

No prediction (3–11 type)

**UNK3–iso2 ...**
63 Seq DWVTRTVEASG Pos 88 98

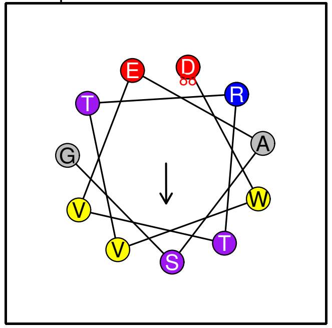

No prediction (3–11 type)

**UNK3–iso1 ...**
64 Seq DWVTRTVEASG Pos 88 98

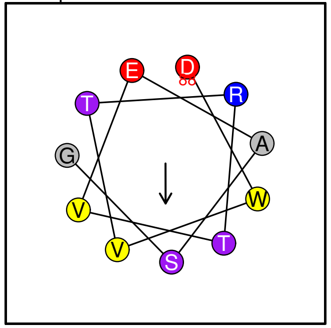

No prediction (3–11 type)

**FBA1–iso3 ...**
65 Seq TDGKKMVDVLV Pos 128 138

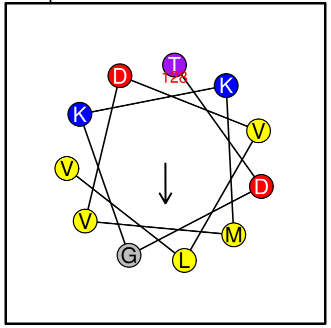

No prediction (3–11 type)

**FBA1–iso3 ...**
66 Seq LDGLASRTAAYYQ Pos 167 179

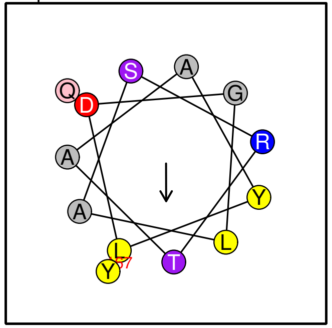

No prediction (3–11 type)

**FBA1–iso3 ...**
67 Seq ANSLAQLGKYT Pos 360 370

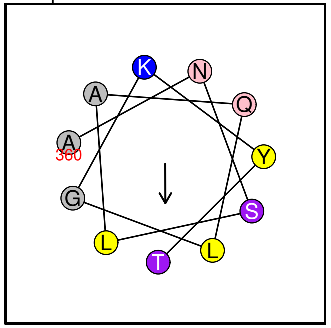

No prediction (3–11 type)

**FBA1–iso2 ...**
68 Seq TDGKKMVDVLV Pos 128 138

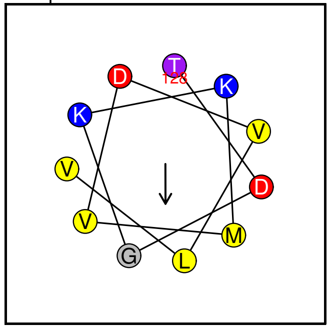

No prediction (3–11 type)

**FBA1–iso2 ...**
69 Seq LDGLASRTAAYYQ Pos 167 179

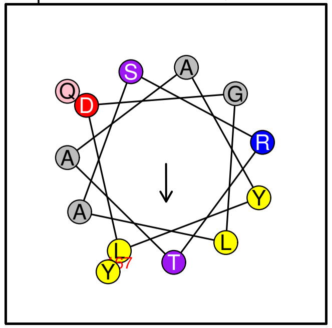

No prediction (3–11 type)

**FBA1–iso1 ...**
70 Seq TDGKKMVDVLV Pos 128 138

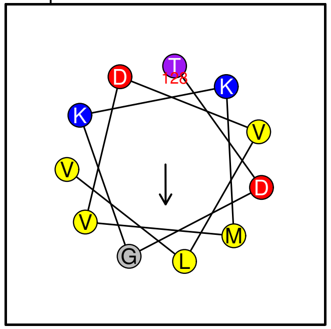

No prediction (3–11 type)

**FBA1–iso1 ...**
71 Seq LDGLASRTAAYYQ Pos 167 179

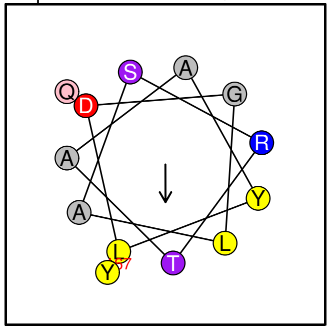

No prediction (3–11 type)

**FBA1–iso1 ...**
72 Seq ANSLAQLGKYT Pos 370 380

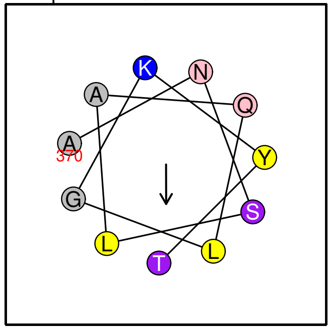

No prediction (3–11 type)

**FBA2−iso1 ...**
73 Seq ANSLAQLGKYT Pos 369 379

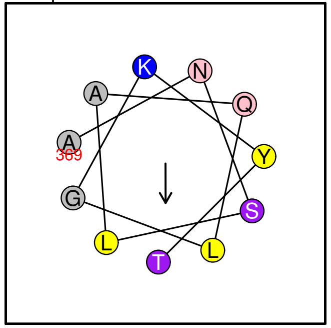

No prediction (3−11 type)

**ABC1K6 ...**
74 Seq AAASLGQVYKG Pos 268 278

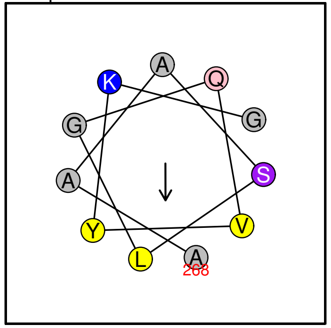

No prediction (3−11 type)

**GMAP−210 ...**
75 Seq AENLEGKVISL Pos 1709 1719

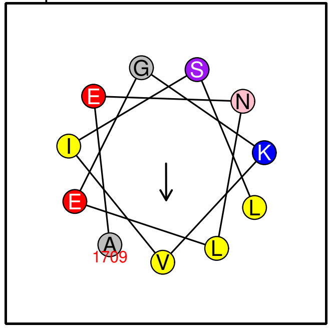

No prediction (3−11 type)

**Est1 ...**
76 Seq GIGGKLKVTVN Pos 47 57

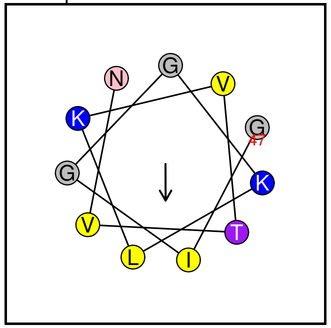

No prediction (3−11 type)

**PEX11 ...**
77 Seq TVGRDKLLRTI Pos 22 32

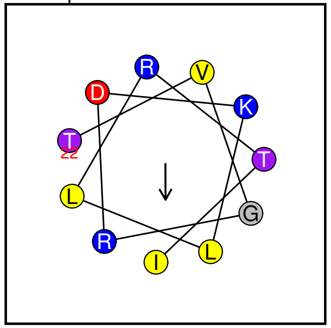

No prediction (3−11 type)

**ABC1K1−iso2 ...**
78 Seq AASLGQVYRAT Pos 244 254

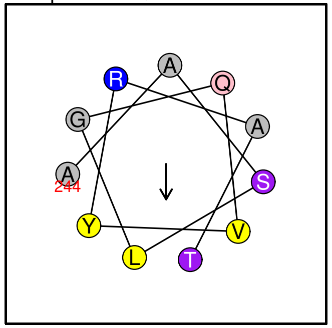

No prediction (3−11 type)

**ABC1K1−iso1 ...**
79 Seq AASLGQVYRAT Pos 244 254

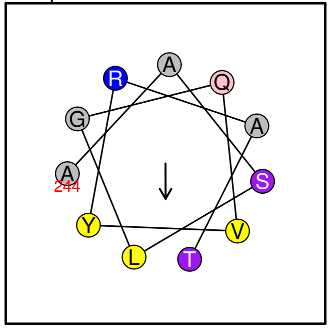

No prediction (3−11 type)

**ABC1K6 ...**
80 Seq DFGLVTKLTDD Pos 437 447

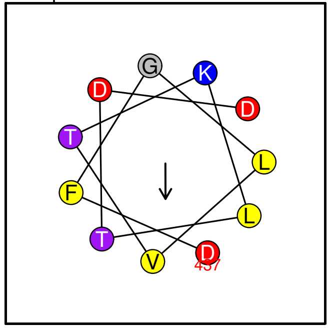

No prediction (3−11 type)

**PES1 ...**
81 Seq KGELAYALDEV Pos 403 413

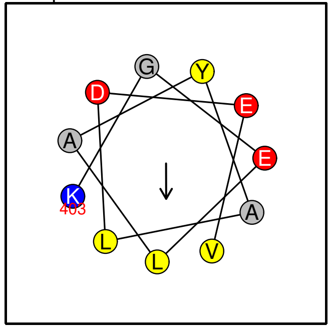

No prediction (3−11 type)

**CCD4 ...**
82 Seq TAARVLTGQYN Pos 219 229

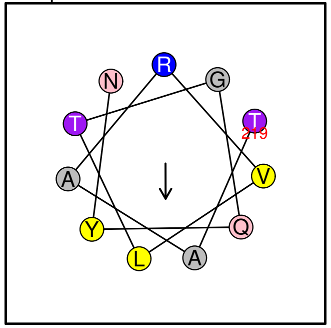

No prediction (3–11 type)

**FBN7b–iso3 ...**
83 Seq EAIYVQLKDAG Pos 45 55

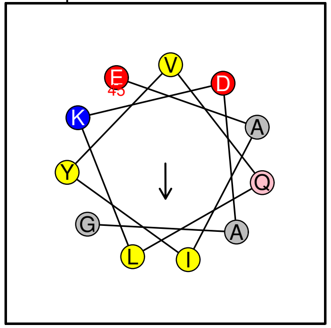

No prediction (3–11 type)

**FBN7b–iso3 ...**
84 Seq QLKSALGQAATTL Pos 192 204

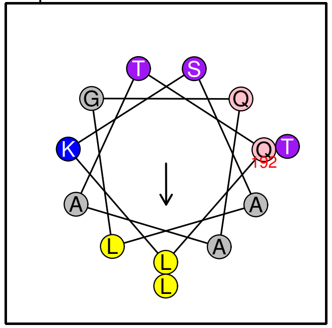

No prediction (3–11 type)

**FBA2–iso2 ...**
85 Seq LDGLSSRTAAY Pos 166 176

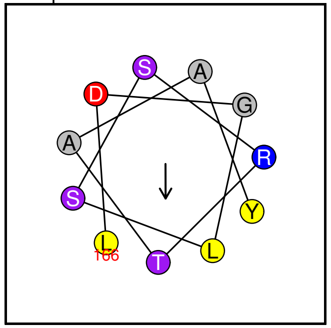

No prediction (3–11 type)

**FBA2–iso1 ...**
86 Seq LDGLSSRTAAY Pos 166 176

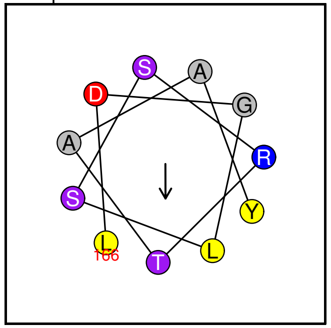

No prediction (3–11 type)

**LOX3 ...**
87 Seq AAGRLKAVLHH Pos 334 344

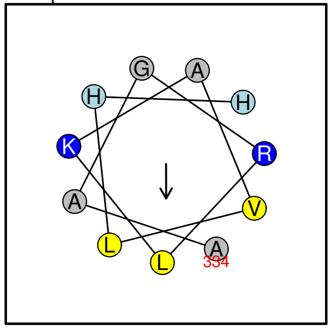

No prediction (3–11 type)

**PLIN3 ...**
88 Seq KGVRTLTAAAV Pos 65 75

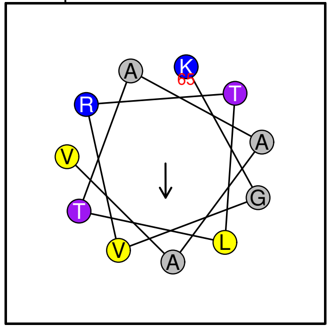

No prediction (3–11 type)

**FBN7b–iso5 ...**
89 Seq QLKSALGQAATTL Pos 125 137

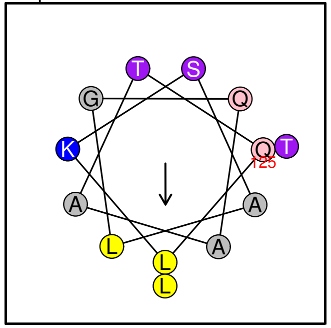

No prediction (3–11 type)

**ABC1K1–iso2 ...**
90 Seq AVVHAVNEDYG Pos 411 421

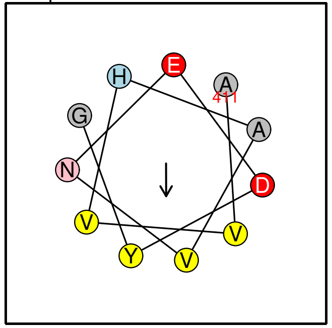

No prediction (3–11 type)

**ABC1K1–iso1 ...**

91 Seq AVVHAVNEDYG Pos 436 446

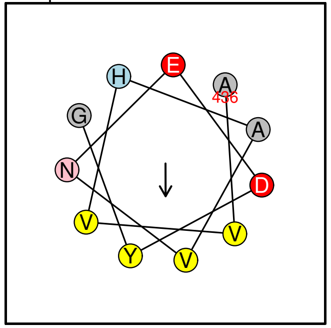

No prediction (3–11 type)

**AOS ...**

92 Seq GEILVEADKLG Pos 294 304

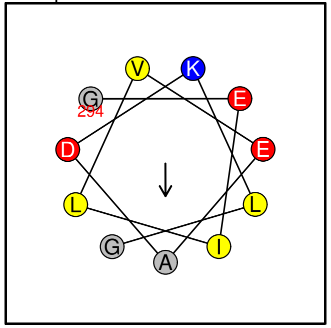

No prediction (3–11 type)

**ABC1K3 ...**

93 Seq VAGGVGQRLAA Pos 683 693

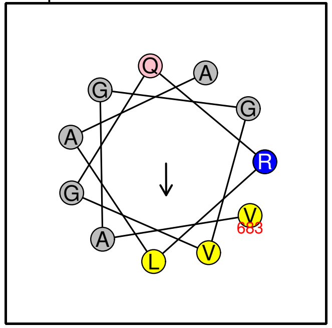

No prediction (3–11 type)

**Unk2–iso2 ...**

94 Seq IGGTLEKVRLN Pos 238 248

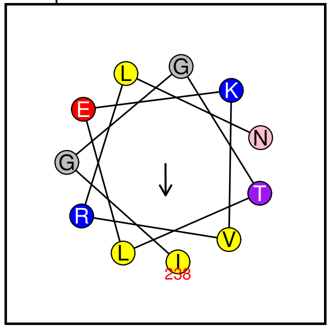

No prediction (3–11 type)

**Unk2–iso1 ...**

95 Seq IGGTLEKVRLN Pos 310 320

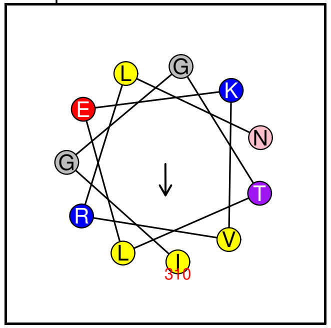

No prediction (3–11 type)

**PLIN3 ...**

96 Seq KSVVTGGVQSV Pos 166 176

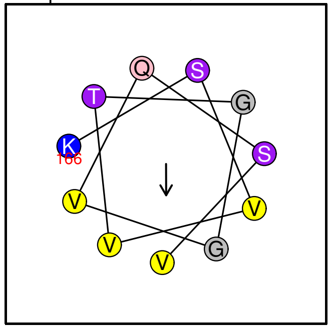

No prediction (3–11 type)

**NDC1 ...**

97 Seq GVELAATISER Pos 257 267

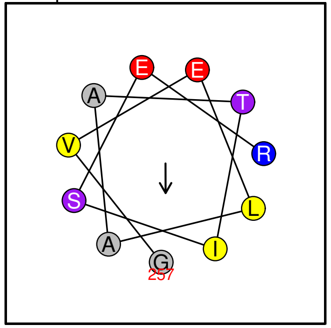

No prediction (3–11 type)

**LOX3 ...**

98 Seq KTKSGVVAAIS Pos 46 56

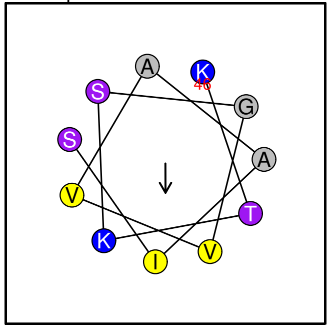

No prediction (3–11 type)

**ABC1K9 ...**

99 Seq EIVVATARGTN Pos 446 456

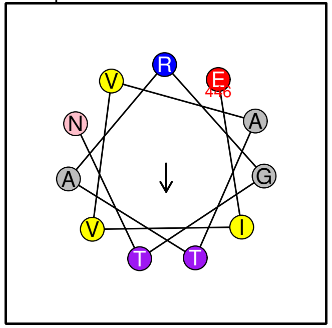

No prediction (3–11 type)

**FBN4 ...**

100 Seq TVKTSGNLSQI Pos 226 236

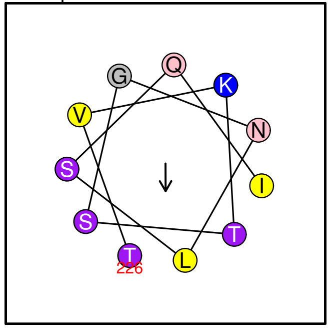


No prediction (3−11 type)

**VTE1 ...**

101 Seq GLTETYENAAL Pos 330 340

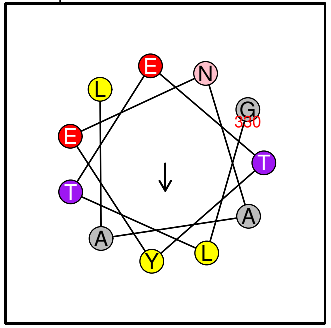


No prediction (3−11 type)

**ABC1K1−iso2 ...**

102 Seq RGAVVSLVSRG Pos 126 136

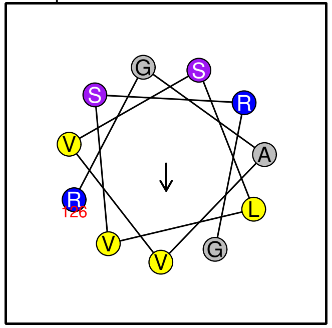


No prediction (3−11 type)

**ABC1K1−iso1 ...**

103 Seq RGAVVSLVSRG Pos 126 136

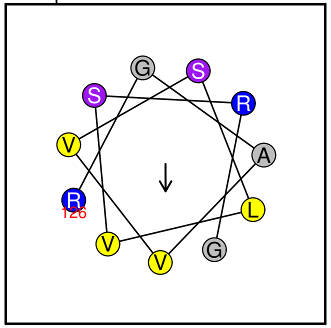


No prediction (3−11 type)
